# Childhood-onset asthma is characterized by airway epithelial hillock-to-squamous differentiation in early life

**DOI:** 10.1101/2023.07.31.549680

**Authors:** Elin T.G. Kersten, J. Patrick Pett, Kristiina Malmström, Yoojin Chun, Marnix R. Jonker, Anna Wilbrey-Clark, Kaylee B Worlock, Maarten van den Berge, Roel C.H. Vermeulen, Judith M. Vonk, Neil Sebire, Jouko Lohi, Wim Timens, Sarah A Teichmann, Supinda Bunyavanich, Marko Z. Nikolić, Martijn C. Nawijn, Mika J. Mäkelä, Kerstin B. Meyer, Gerard H. Koppelman

## Abstract

Childhood-onset asthma is characterized by Type 2-inflammation and airway wall remodeling, but mechanisms of asthma development in the first years of life remain unclear. Here, we investigate transcriptional changes in airway wall biopsies of 22 symptomatic one year old children and relate these to asthma at school age. We demonstrate that pre-asthmatic children (n = 10) overexpressed a gene signature characteristic for an airway epithelial differentiation trajectory via hillock cells towards squamous cells (adjusted p-value 8.06e-16), whilst there was no association with gene signatures of Type 2-inflammation or eosinophil activation. Genes expressed along this trajectory are linked to an altered epithelial barrier function, innate immune activation and extracellular matrix remodeling. Functional GWAS analysis supports a causal link between childhood-onset, but not adult-onset asthma, and the hillock-squamous cell differentiation trajectory. Next, we confirmed the presence of hillock-like cells at the RNA and protein level in pediatric upper and lower airway samples. These findings identify a novel mechanism by which an aberrant airway epithelial differentiation trajectory may contribute to a pre-asthmatic state, highlighting the difference between the early origins of childhood-onset asthma and adult asthma, and point to possible new targets for the early diagnosis and treatment of asthma in the first two years of life.

**One Sentence Summary:** RNA sequencing in bronchial biopsies from wheezing infants and children < 2 years shows evidence for an airway epithelial hillock-to-squamous differentiation pathway that marks the development of asthma.

## INTRODUCTION

Asthma is a highly prevalent respiratory disease that often has its onset in early childhood. It is the outcome of the interaction of genetic susceptibility with environmental exposures, including viral infections and air pollutants, leading to chronic disease (*1*). While research in adults suggests that persistent asthma involves eosinophilic, Type 2 airway inflammation and remodeling of the airway wall (*2*), very little is known about early changes in the airways of preschool children developing asthma (*3, 4*). In symptomatic, wheezing preschool children that do not yet meet the criteria to diagnose asthma, but go on to develop asthma at school age; a group we term “pre-asthmatic”, alterations in the airway wall may already be present. Only four cohorts have combined investigations of bronchial biopsies in early life with follow-up to a clinical diagnosis of asthma at school age (*5–8*). At 2-3 years of age, features of remodeling, including increased reticular basement membrane (RBM) thickness (*7*) and airway smooth muscle (ASM) hyperplasia (*6*), rather than markers of Type 2-inflammation (*8*), predict progression to asthma at school age. In an even younger cohort of 53 corticosteroid-naive children <2 years of age with recurrent severe lower respiratory tract symptoms including wheeze, we previously reported no correlation of RBM thickness, ASM fraction, or endobronchial inflammatory cell counts with persistent asthma at school age (*5*). Thus, structural changes in the airway wall may precede airway inflammation in pre-asthmatic children, but potential mechanisms that govern this are unknown.

Childhood-onset asthma has a strong genetic contribution, with a distinct set of genes involved compared to adult-onset disease (*9, 10*), suggesting that specific molecular mechanisms underlie disease inception in childhood. Here, we performed bulk RNA-sequencing (bulkRNA-seq) on airway wall biopsies from children <2 years with severe wheeze, who subsequently either do or do not develop asthma, allowing us to identify gene expression changes and enriched pathways that are linked to a pre-asthmatic disease state. Next, we used single cell RNA sequencing (scRNA-seq) data from a separate study that profiled lower airway samples from healthy preschool children to study which cell types and states express the pre-asthmatic gene signature. Finally, we utilised summary statistics from genome wide association studies (GWAS) of both childhood- and adult onset asthma (*11, 12*) to test whether single nucleotide polymorphisms (SNPs) associated with childhood-onset asthma can be linked to the cell-types that express the pre-asthmatic gene signature.

## RESULTS

### A gene expression signature of squamous cell differentiation is associated with childhood onset asthma

To investigate which transcriptional programs in the airway wall of symptomatic preschool children are associated with asthma at school age, we performed bulkRNA-seq of archived bronchial biopsies of children <2 years of age with severe recurrent respiratory symptoms as described previously (*5*) and grouped them into “asthma” and “transient severe wheeze” based on the outcomes of their extensive clinical follow-up at 6-8 years. Asthma at follow-up was defined as physician-diagnosed asthma with regular or intermittent use of inhaled corticosteroids (ICS) or leukotriene antagonists during the past year. After quality control, twenty-two samples were retained (see online supplement). Of these, 10/22 (45%) children had asthma and 12/22 (55%) had no asthma at age 6-8 years. Clinical characteristics at time of biopsy and follow-up are shown in Table 1, whereas previous histological characterization of these biopsies (*13*) is summarized in Supplementary Table S1.

**Table 1.** Patient characteristics and clinical outcomes at school age.

|  | **Asthma (n=10)** | **Transient wheeze (n=12)** |
| --- | --- | --- |
| **Baseline information at 1y** |  |  |
| Gender, male | 9 (90%) | 11 (92%) |
| Age (months) | 13.6 ± 7.3 | 13.5 ± 6.3 |
| Height (cm) | 76.1 ± 9.1 | 74.5 ± 5.4 |
| Weight (kg) | 10.1 ± 2.6 | 9.9 ± 1.3 |
| Duration of symptoms (months) | 7.5 ± 5.1 | 8.9 ± 4.6 |
| Wheezing apart from colds | 6 (60%) | 10 (83%) |
| Atopic dermatitis | 2 (20%) | 2 (17%) |
| FRC (z-score) | 0.5 ± 1.4 | 0.5 ± 1.2 |
| Skin prick test + | 3 (30%) | 2 (17%) |
| Total IgE (kU/L) | 68.5 ± 96.3 | 37.8 ± 51.6 |
| Eosinophils (%) | 2.5 ± 1.0 | 3.1 ± 1.9 |
| FeNO (ppb) | 19.7 ± 15.6 | 16.9 ± 6.2 |
| Parental smoking | 3 (30%) | 6 (50%) |
| Parental asthma | 5 (50%) | 6 (50%) |
| Parental allergy | 9 (90%) | 10 (83%) |
| **Follow up** |  |  |
| Age follow up (years) | 7.0 ± 1.7 | 7.9 ± 1.5 |
| Height (cm) | 123.8 ± 11.9 | 126.3 ± 7.8 |
| Weight (kg) | 28.3 ± 9.4 | 25.9 ± 4.0 |
| FeNO (ppb) | 12.0 ± 11.1 | 8.8 ± 5.7 |
| Airway hyperresponsiveness | 8 (80%) | 7 (58%) |
| Atopic dermatitis | 1 (10%) | 5 (42%) |
| Atopy | 3 (30%) | 8 (67%) |
| Eosinophils (%) | 5.7 ± 5.0 | 6.3 ± 7.6 |

Legend: All values are represented as mean ( ± SD) or number (percentage). Asthma was defined as physician-diagnosed asthma ever and regular or intermittent use of ICSs or LTRAs during the past year. AHR was defined as a PD<sub>40</sub> of 0.4 mg or less on a methacholine challenge test. Atopy was defined by a positive skin prick test result to food or aeroallergens or presence of atopic dermatitis.

We did not observe individual differentially expressed (DE) genes at genome-wide significance between asthma and transient wheeze, and between the presence or absence of atopy and airway hyperresponsiveness (AHR) at follow up (Tables S2-S4). To test whether gene programs belonging to a specific biological pathway were DE between asthma and transient wheeze, we next performed pathway and gene ontology (GO) overrepresentation analysis on all nominally significant DE genes (p < 0.05, n=404, Table S2). We identified upregulated pathways and GO terms enriched in the early life biopsies from pre-asthmatic children, including “keratinization”, “epidermis development” and “cell cycle” pathways (Fig. 1A, C and Table S5, S6).

**Fig. 1.**
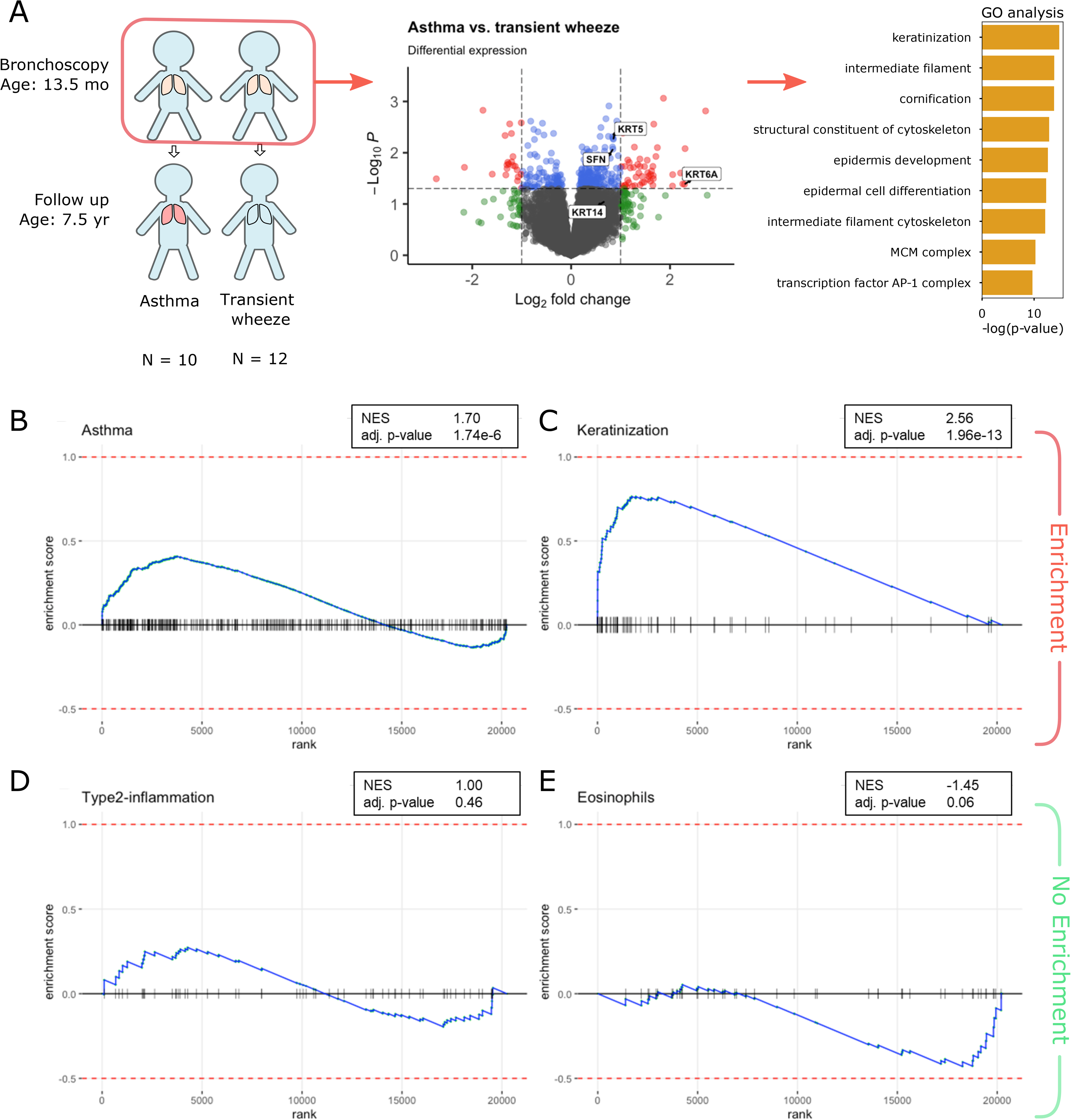
Study design and main result of DEA. **(A)** Study design of DEA between children < 2 years with asthma at school age and transient severe wheeze. Middle panel shows the results from DEA with cutoffs for (nominal) p-value at 0.05 used for selecting the geneset for GO analysis, and the cutoff for log2fold difference at 0.8 used for the pre-asthma gene signature. Right panel shows the significant results from the GO overrepresentation analysis. **(B-E)** Enrichment plots showing the normalized enrichment scores with adjusted p-value of selected gene signatures in children <2 years in relation to asthma at school age. **(B)** enrichment plot of genes upregulated in persistent asthma (*14*) **(C)** enrichment plot of genes in keratinization GO set **(D)** enrichment plot of genes upregulated in Type2-inflammation (*15*) **(E)** enrichment plot of genes in geneset of activated airway eosinophils (*16*)

### Asthma associated gene expression in pre-asthmatic children is not driven by eosinophilia or Type 2-inflammation

Next, we investigated whether gene expression patterns characteristic of asthma in adults and older children can already be observed in airway wall biopsies in pre-asthmatic children. We therefore evaluated enrichment of three previously reported gene sets: (1) genes expressed in airway samples associated with asthma in a large meta-analysis (*14*), (2) genes characteristic for Type 2-inflammation (*15*) and (3) genes expressed in activated airway eosinophils (*16*). The asthma gene set was significantly enriched in the early life biopsies from pre-asthmatic children compared to transient wheeze (normalized enrichment score (NES) 1.70, adj. p-value 1.74e-6; Fig. 1B, Table S7), yet this was not driven by genes associated with Type2-inflammation (Fig. 1D, Table S8) or activation of eosinophils (Fig. 1E, Tables S9), but by genes relating to epithelial cell proliferation and differentiation (Table S7). Thus, these early life biopsies did not show transcriptomic evidence of eosinophilic or Type2-inflammation, yet appear to be characterized by a specific epithelial state.

### Projection of single-cell data links hillock and squamous cells to pre-asthma

To identify whether the transcriptomic differences observed between biopsies from pre-asthmatic children and those with transient wheeze reflect changes in cell type composition of the airway wall, we utilized a recently published airway cell atlas as a reference dataset (*17*). From this dataset we selected tracheal brush samples of 9 healthy donors below 4 years of age (9551 cells in total) for all further analyses (Fig. 2A), as these most closely matched our study population.

**Fig. 2.**
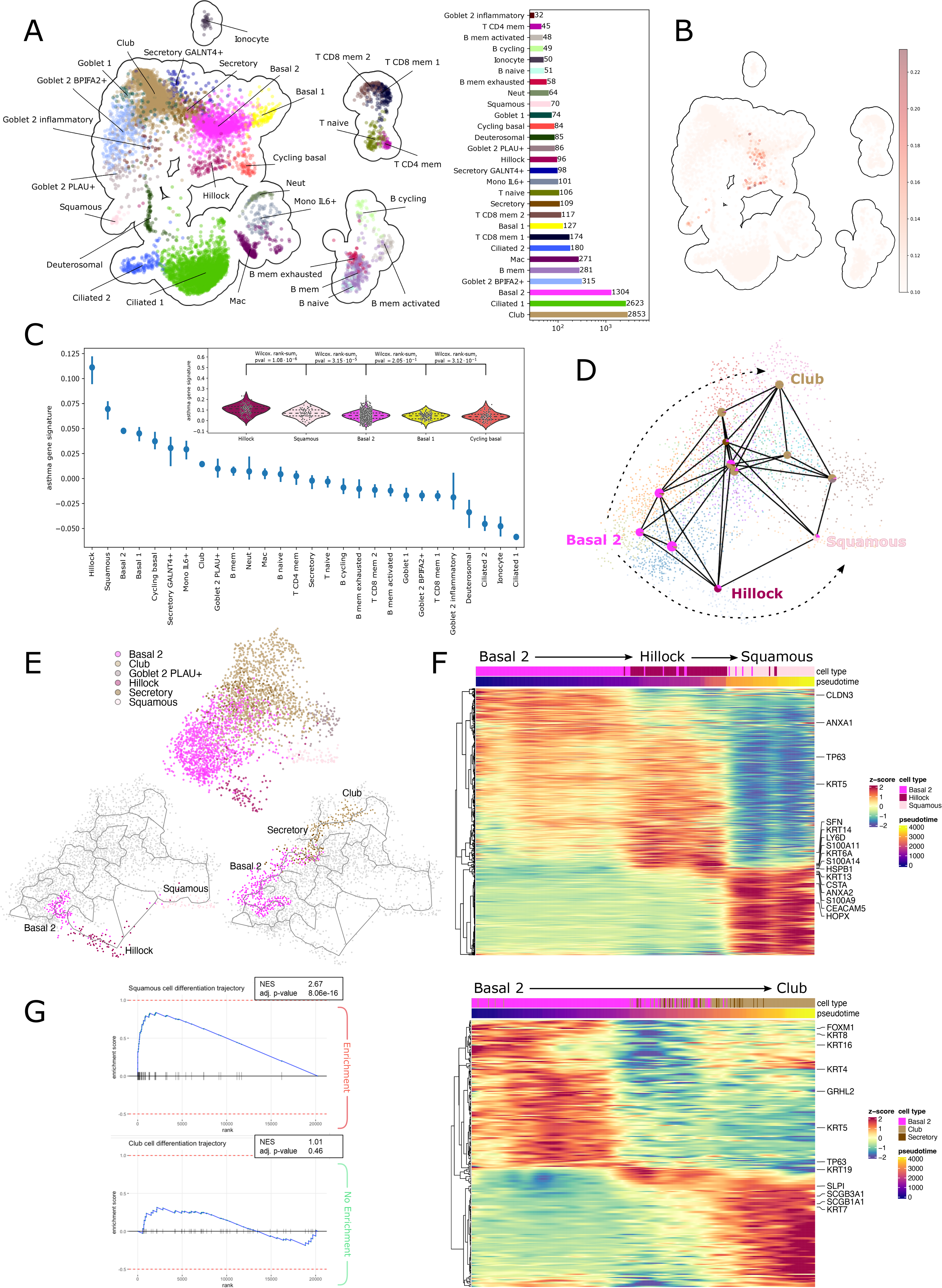
Single cell data links pre-asthma to hillock and squamous cells. **(A)** UMAP with cell type annotations showing tracheal samples of healthy donors below 4 years of age. The number of cells per cell type is shown on the right. **(B)** Gene expression of pre-asthma signature (DE cutoff: p-value<0.05 and LFD>0.8) in UMAP. Cell-level scores were computed for the gene signature. **(C)** Ranking of cell subsets based on the median score (points) with 95% confidence intervals (lines). The inset-plot shows the distributions of the top 5 cell subsets with hillock cells at the first position. Also shown are p-values for wilcoxon rank-sum tests between successive positions. **(D)** PAGA of leiden clusters showing connectivities between Basal 2, Hillock, Squamous, Club, Secretory and Goblet 2 PLAU+ cell subsets. **(E)** Trajectories selected using Monocle3 principle graph. Bottom left: Basal 2 → Hillock →Squamous trajectory subsetted from top panel. Bottom right: Basal 2 → Club trajectory. **(F)** Heatmaps showing the expression of DE genes along the two trajectories. Selected genes are marked on the right. Differential expression was tested using a spatial autocorrelation test. **(G)** Enrichment plots of squamous differentiation trajectory and club differentiation trajectory gene signatures showing the normalized enrichment scores with adjusted p-value in children < 2 years in relation to asthma at school age.

First, we analyzed the expression of a composite pre-asthma gene signature, encompassing the nominal early life pre-asthma associated DE gene list (p-value < 0.05, Table S2). We compared the relative expression of the signature across all cell types in the healthy, pediatric scRNA-seq dataset using the original labels (*17*) (Fig. 2B,C). Interestingly, the cluster epithelial cells annotated as hillock cells consistently showed highest expression, followed by squamous and basal 2 (suprabasal) cell subsets. Indeed, the pre-asthma gene signature includes the cytokeratin genes *KRT13* and *KRT5*, recently described as marker genes of hillock cells in mice (*18*).

These findings suggested that gene expression of pre-asthma was driven by specific airway epithelial cell subsets. To test this hypothesis, we computed marker gene lists for all 14 annotated epithelial cell subsets in the single cell dataset using a logistic regression approach (Table S10). The top-50 marker genes per epithelial cell type were obtained with generally small overlap and a good distinction between labels (Fig. S1A). We then performed gene set enrichment analysis (GSEA) with these top-50 marker genes per label in our bulkRNA-seq dataset. The top-50 marker genes for squamous, hillock, cycling hillock, cycling basal and goblet *PLAU*+ cells were enriched in pre-asthmatic children compared to transient wheeze (adjusted p-value <0.005; Table S11, Fig. S2), suggesting that presence and/or activation of these cells characterized pre-asthma.

### A distinct differentiation trajectory towards squamous cells is linked to pre-asthma

Squamous cells differentiate from basal cells in the airway epithelium through an alternate trajectory to the basal-club-goblet cell or basal-club-ciliated cell trajectories (*19*). Murine hillock cells have previously been described as an intermediate transitory cell state in squamous epithelial differentiation in the trachea (*18*). We therefore investigated whether the pre-asthma-associated epithelial cell subsets constitute a distinct differentiation trajectory. In our scRNA-seq dataset, we applied PAGA analysis to infer potential differentiation trajectories in healthy pediatric tracheal epithelium (*20*). We observed that the different epithelial cell subsets formed a continuum lacking discrete cell types, underscoring the plasticity of the airway epithelium, which is also reflected by the large connectivity between basal 2, hillock, squamous, club, secretory and goblet 2 *PLAU*+ cells (Fig. 2D). The PAGA graph linked hillock to squamous cell subsets, with a portion of hillock cells placed on a predicted transition between hillock and squamous cells.

We next applied Monocle3 (*21, 22*) to select two distinct differentiation trajectories starting from basal 2 cells: (I) one connecting to club cells, and (II) one to squamous via hillock cells (Fig. 2E). Differential gene-expression along the basal 2-hillock-squamous trajectory highlighted many hillock specific marker genes appearing close to the transition to squamous cells including *SFN*, *KRT14* and *S100A11*, with several other markers like *KRT13*, *ANXA2* and *S100A9* being expressed in both cell states (Fig. 2F top). In contrast, luminal cytokeratin *KRT19* and secreted proteins like *SLPI* that protect the epithelial surface are expressed at the transition towards club cells (Fig. 2F bottom). Taken together, these results indicate that the pre-asthmatic gene signature reflects an airway epithelial differentiation trajectory from basal 2, via hillock, towards squamous cells, rather than a single epithelial cell state. To formally test this, we generated a gene signature with the top 50 genes differentially expressed in the two selected differentiation trajectories (Table S12). GSEA in the bulkRNA-seq dataset with these two trajectory-specific gene signatures showed strong enrichment for the squamous cell differentiation gene signature (NES 2.67, adjusted p-value 8.06e-16), but not for the club cell differentiation gene signature (NES 1.01, adjusted p-value 0.46) (Fig. 2G) in the early life biopsies from pre-asthmatic children compared to transient wheeze.

To the best of our knowledge, there is no other bronchial biopsy RNAseq dataset from young pre-asthmatic children available for replication of our results. We therefore aimed to replicate our findings in two independent nasal brush bulkRNA-seq datasets of adolescents with mild and severe asthma. From the scRNA-seq dataset, we now selected a subset of nasal samples of healthy donors below 4 years of age and computed marker gene signatures of the squamous and club cell differentiation trajectories using logistic regression. We performed GSEA with these gene signatures in the Prevention and Incidence of Asthma and Mite Allergy (PIAMA) birth cohort (*23*), consisting of 326 adolescents including 24 with mild asthma and the Children from the AiRway In Asthma (ARIA) study cohort (*24*), consisting of 34 children with severe persistent asthma and 122 healthy controls (see online supplement for details). GSEA with these two trajectory gene signatures showed downregulation of the squamous cell differentiation gene signature (NES -1.39, adj. p-value 0.10 in PIAMA and NES -1.55, adj. p-value 0.04 in ARIA) and an inconsistent pattern of the club cell differentiation gene signature (NES -0.69, adj. p-value 0.92 in PIAMA and NES 1.60, adj. p-value 0.02 in ARIA) in adolescents with asthma (Fig. S3A-D). We note that these cohorts differ from our discovery cohort in the sampling locations (nasal brush samples versus bronchial biopsies) as well as the average age of the participants (PIAMA mean 16.3 ± 0.2 years, ARIA mean 13.1 ± 3.7 years).

### GWAS SNPs link childhood-onset asthma genes to hillock cell transcriptomes

We hypothesized that genes relating to childhood-onset asthma are preferentially expressed in airway cell-types linked to pre-asthma. Therefore, we performed a functional GWAS (fGWAS) analysis (*25, 26*) which tests for significant enrichment of disease-associated variants per cell type in corresponding regions of transcribed genes, taking into account both strongly and weakly associated SNPs by using the full GWAS summary statistics (*9, 10*) (Fig. 3A).

**Fig. 3.**
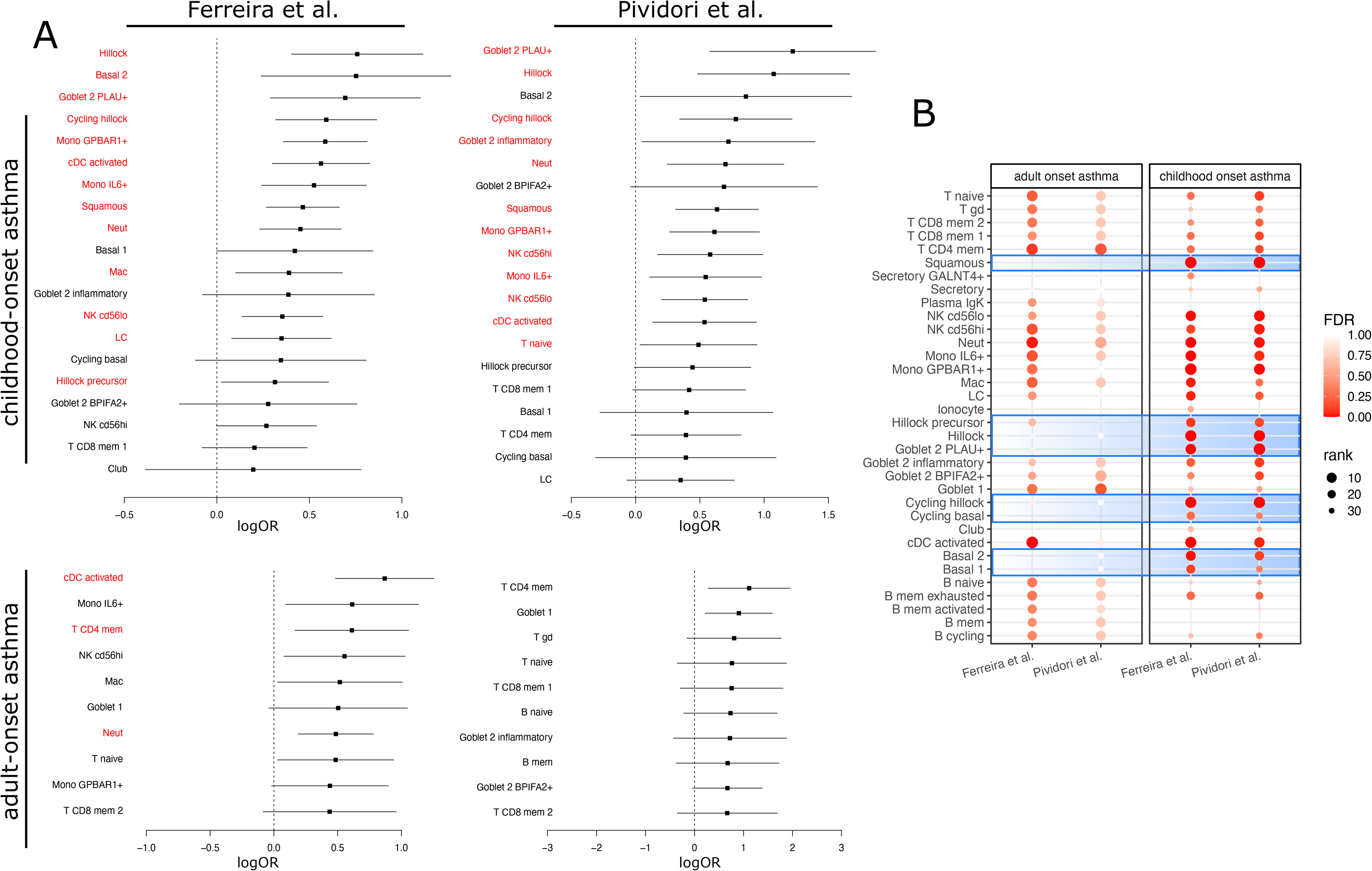
Functional GWAS results. **(A)** Forest plots for four sets of summary statistics. The logOR is shown on the x-axes and significant enrichments are marked in red. Cell subsets are ranked by effect size. The top 20 and top 10 cell subsets are shown for childhood-onset and adult-onset asthma, respectively. **(B)** Dot plot comparing the rankings of cell subsets across studies. Different subsets of hillock cells and basal cells as well as squamous and goblet PLAU+ cells show the largest difference between childhood- and adult-onset asthma.

Strikingly, fGWAS using the childhood-onset asthma SNPs identified hillock cells as the top enriched cell subset in our scRNA-seq dataset of healthy, early life tracheal brushings. In addition, goblet *PLAU*+ cells, an inflammatory type of goblet cells, and basal 2 (suprabasal) cells also showed strong enrichment; the latter result was only significant for the Ferreira *et al.* dataset. Squamous cells were also significantly enriched, but ranked lower in the list of cell subsets. Notably, various immune cells typically linked to asthma did not show enrichment in the fGWAS analysis for childhood onset asthma. We also analyzed the contribution of genes with the largest disease association, which showed consistent results (Fig. S4).

In contrast to our findings for childhood-onset asthma, fGWAS analysis with adult-onset asthma summary statistics identified a number of immune cell subsets as the most enriched cell types (Fig. 3A). Remarkably, hillock cells and goblet *PLAU+* cells were not significantly enriched in the adult-onset asthma fGWAS. Overall, the largest differences were observed between childhood- and adult-onset asthma for hillock-like (hillock, cycling hillock, hillock precursor), squamous, basal (basal1, basal2, cycling basal) and goblet *PLAU+* cell subsets (Fig. 3B).

### Protein stainings and RNAscope confirm the presence of cells expressing squamous differentiation trajectory markers

Hillock cells were first described as anatomically distinct hillock-shaped clusters of epithelial cells in the mouse trachea (*18*), and scRNA-seq of pediatric airway brushings provided evidence for their presence in the pediatric human airway wall (*17*), but this has not been confirmed by spatial methods. To confirm the presence of epithelial cells co-expressing hillock markers in the airway wall in humans, we performed single molecule *in situ* hybridization (smFISH) using commercial RNAscope probes for *KRT14* and *KRT6A*. This analysis showed the presence of *KRT14/KRT6A* co-expressing cells, consistent with hillock cells, and their presence was observed both in the nose (pediatric samples) and lower airways (adult samples) (Fig. S5). We observed patches of cells with no apparent “hillock” morphology as described for mice (*18*).

We subsequently performed immunofluorescence stainings with KRT5, a basal cell marker, and KRT14, a squamous differentiation trajectory marker, in archived paraffin embedded airway wall biopsies (n = 36) from the trachea of the severely wheezing children, in a subset of whom we also performed the bulkRNA-seq analysis. Here, we could confirm the presence of cells co-expressing KRT5 and KRT14 (Fig. 4), with similar patterns of double-positive cells as were observed in the RNAscope analysis. Two independent, blinded observers scored the presence of KRT5/KRT14 co-expression; however no significant predictive association was found of the number of KRT5/KRT14 double-positive cells with the outcome of asthma at 6-8 years (Chi-square test = 2.59, p-value = 0.11).

**Fig. 4.**
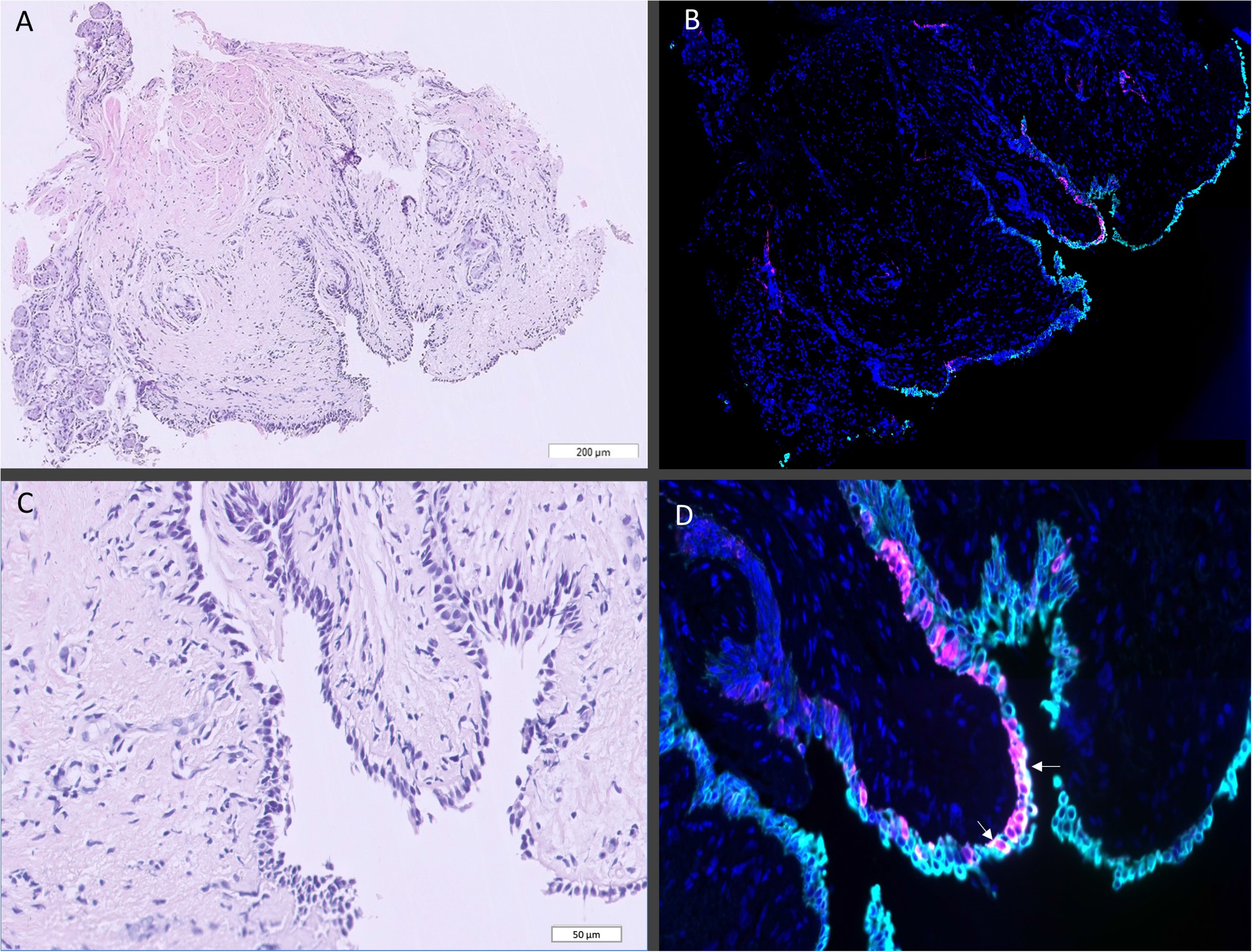
H&E and immunofluorescent staining. Staining with cytokeratin 5 (cyan) and cytokeratin 14 (magenta) with DAPI (dark blue) stained nuclei of bronchial biopsy from a 6 month old boy with severe recurrent wheeze. Double positive cells are white and appear in patches along the epithelial layer (arrow). (A) H&E staining of whole biopsy. (B) Immunofluorescent staining of whole biopsy. (C) Zoomed in H&E staining. (D) Cytokeratin 5 and 14 double staining, zoomed in on airway epithelial cells with double positive cells (white, arrow).

## DISCUSSION

This study is the first to implicate a bronchial epithelial hillock-to-squamous differentiation trajectory in the onset of childhood asthma in the first years of life. We provide evidence for the hillock-to-squamous trajectory in the airway epithelium in early life at several levels: We define a pre-asthma associated gene signature that is overexpressed in airway epithelial cells along this trajectory. We find that cell-type specific genes for hillock-to-squamous epithelial cell subsets are enriched for SNPs linked to childhood-onset asthma risk, while cell-type specific genes of immune cell subsets are enriched for SNPs linked to adult-onset asthma. In addition, we provide transcriptional evidence that established mechanisms of persistent asthma, including Type 2-inflammation and airway eosinophilia, are not enriched in wheezing, pre-asthmatic children. Finally, we confirm the presence of hillock-like cells in the upper and lower airways of children at the RNA and protein level.

In our study, we describe a specific transitional epithelial cell state mapping to the hillock-to-squamous differentiation trajectory, which is strongly enriched for childhood-onset asthma genes. Various studies have highlighted phenotypic (*27*) and genetic (*9, 10*) differences between childhood- and adult-onset asthma. The identification of this alternative epithelial differentiation trajectory may explain many previous observations in childhood-onset asthma, as this pathway may be linked to (1) altered epithelial barrier function, (2) immune activation and (3) remodeling of the extracellular matrix (ECM).

First, the hillock-to-squamous differentiation may be linked to reduced epithelial barrier function. Squamous epithelium is a stratified epithelial cell surface phenotype usually found a.o. in the skin and esophagus (*28*). However, a recent study showed the presence of patches exhibiting squamous differentiation in healthy human trachea, enriched with hillock and squamous gene expression such as *KRT6a* and *KRT13* (*28*). In air-liquid interface (ALI) cultures, squamous epithelial cells showed epithelial unjamming and compromised epithelial barrier integrity (*28*). Genetic studies of asthma, atopic dermatitis (AD) and allergic rhinitis have indicated a very strong overlap of causal genes of these allergic diseases, specifically when age of onset was taken into account (*29, 30*). Many AD genes are located in the epidermal differentiation complex at chromosome 1q, that includes genes important for keratinocyte differentiation and cornification, S100 calcium-binding and S100 Fused type proteins, such as filaggrin. Interestingly, AD skin transcriptomics show a remarkable overlap with the squamous differentiation trajectory gene signature, with upregulation of *KRT13, KRT6a, S100A8, S100A9* and *SPRR3* (*31*). Thus, the link between the hillock-to-squamous differentiation pathway and an altered epithelial barrier function may have a genetic origin that is shared with other allergic diseases.

Second, cells differentiating towards squamous cells may influence the immune response through upregulation of proteins such as innate cytokines, S100 calcium-binding proteins or annexins. Squamous epithelial cells have been shown *in vitro* to express several inflammatory cytokines and chemokines, displaying an upregulation of multiple inflammatory pathways such as IL-6-JAK-STAT3 signaling (*28*). S100A8 and S100A9, which are upregulated along the hillock-to-squamous trajectory (Fig. S1B), are damage associated proteins that induce the release of innate pro-inflammatory cytokines, such as IL-6 in airway epithelial cells (*32*). In an *in vitro* model of primary airway epithelial cells, rhinovirus infection and its sequelae, including expression of innate inflammatory cytokines and release of IL-6, were much more pronounced in a culture that had resulted in squamous epithelium compared to a culture of differentiated pseudostratified epithelium (*33*). Other genes expressed along the hillock-to-squamous trajectory, such as annexins and cystatins, are known for their roles in innate immune modulation. For example, ANXA2 interferes with arachidonic acid release and eicosanoid formation (*34*), biologically active molecules that play an important role in Type2-inflammation. SNPs in the *ANXA1* gene have recently been associated with early-onset persistent wheeze (*35*). This implies that the hillock-to-squamous differentiation pathway may be related to an altered immune response to epithelial damage.

Third, interaction between epithelial and mesenchymal cells has been proposed to be central in asthma development (*36*). ECM remodeling may occur by secretion of proteins like stratifin (SFN, 14-3-3σ) by epithelial cells along the hillock-to-squamous trajectory (Fig. S1B). Stratifin has been linked to the regulation of proliferation, cell adhesion and ECM remodeling (*37, 38*). Releasable 14-3-3 proteins from airway epithelial cells can induce MMP-1 in lung fibroblasts (*39*), which may enhance ASM hyperplasia and airway obstruction in asthma (*40*). Thus, this may provide a possible mechanism of ASM hyperplasia and thickening of the RBM occurring in pre-asthma without the presence of Type 2-inflammation (*6*).

Transcriptomic signatures of Type 2-inflammation and eosinophilia were not enriched in pre-asthmatic children compared to transient wheeze. This confirms our previous findings from histology (Table S1) (*5*) and adds to the growing body of evidence that pre-asthma is not characterized by eosinophilia or Type 2-inflammation.

We propose that not one specific cell type, but a dysregulated epithelial differentiation trajectory across several cell types and states (from basal 2, through hillock, goblet 2 PLAU+ and squamous cells) is characteristic of pre-asthma. This may explain why the transcriptomic signature of this differentiation trajectory was enriched in pre-asthma, but not the number of hillock cells as detected by protein stainings. Interestingly, we observed that hillock cells appear to group in suprabasal layers, but we do not observe the typical “hillock” structure as described by Montoro et al. in the mouse (*18*). This is consistent with previous observations of relatively solitary hillock cells that were KRT4/KRT13 double positive in nasal tissue (*41*).

A critical question is what drives the differentiation of basal epithelial cells. We propose that epithelial injury due to recurrent viral infections in these severely wheezing children, induced a repair response. In at-risk infants, this repair response involves a proliferative response, with subsequent activation of the epithelial differentiation complex genes and transitioning of the basal epithelial cells into a hillock-to-squamous trajectory. Recently, Zhang et al showed that Wnt/β-catenin signaling pathway activation serves as a driving force in promoting squamous cell fate from airway basal cells (*28*). In contrast, in not-at-risk infants, the airway epithelial cells might use repopulation of basal cells and restoration of regular differentiation pathways as a repair response to deal with epithelial injury. We speculate that the genetic make-up predisposing to asthma is expressed as a higher tendency to enter a squamous cell differentiation trajectory as part of the epithelial repair response. The genes that are expressed along this differentiation trajectory, and the respective roles of their translated proteins, such as S100 proteins, annexins, cystatins and stratifin, warrant further functional investigation as potential novel therapeutic targets or diagnostic biomarkers.

The results of this study should be interpreted carefully, as our study has some limitations. First of all, the sample size of 22 is small, which likely explains why there were no genes DE at genome-wide significance. The final outcome, current asthma at school age, is not as straightforward as it may seem; a child’s phenotype and the presence of asthma symptoms may vary over time. Childhood-onset asthma also covers several phenotypes and the patient selection based on severe recurrent respiratory symptoms might select a specific subgroup of asthma patients who are more susceptible to recurrent viral infections. Therefore, our results need to be replicated in an independent cohort that combines RNA-seq of early life bronchial biopsies with follow-up at school age. However, to the best of our knowledge, no such cohort is currently available. We therefore aimed to replicate our findings in two nasal brush RNA-seq datasets of adolescents with mild and severe asthma. Interestingly, we didn’t find enrichment for the squamous differentiation trajectory signature in these cohorts. This could be due to the differences in age, location, sampling method or medication use. The nasal epithelium is derived from a different embryonic origin and scRNA-seq has shown that cell type proportions vary between nasal brushings and tracheal biopsies (*41*) with overall higher expression of squamous differentiation trajectory genes in the nasal epithelium. In the currently available data, it is impossible to distinguish whether the relative upregulation of a gene signature is due to upregulation of certain cell types or states, or an increased expression of certain genes in all cell types.

In conclusion, we identified a distinct pattern of hillock-to-squamous differentiation in the early development of asthma, without evidence of Type 2-high, eosinophilic inflammation. We hypothesize that the transitional hillock cells and squamous epithelial cells may contribute to subsequent ECM remodeling and an innate immune response, creating a microenvironment that facilitates the development of asthma. Our data define a pre-asthma state that is different in early life, yet predisposes for chronic, persistent asthma.

## MATERIALS AND METHODS

### Study design

The study population consisted of 49 Caucasian infants referred to Helsinki University Central Hospital for early onset doctor diagnosed wheeze and recurrent troublesome breathing. Additional details are in the online supplement. For 26 children from this cohort, an additional biopsy was available for bulkRNA-seq.

At school age, children were invited to participate in a follow-up study. The child’s parent/guardian filled in a questionnaire on asthma symptoms and medication. Current asthma was defined as physician-diagnosed asthma and regular or intermittent use of ICS or leukotriene antagonists during the past year. Atopy was defined by a positive skin prick test result to food or aeroallergens. Lung function and bronchodilator responsiveness were measured using a pneumotachograph-based flow-volume spirometer (Masterscreen Pneumo, Jaeger GmbH, Wurzburg, Germany). AHR was evaluated with a dosimetric methacholine challenge. Written informed consent was obtained from the children and one of the parents. The Ethics Committee of Diseases of Children and Adolescents at the Helsinki University Central Hospital approved the study protocol.

### Sample collection

Bronchoscopy was performed on clinical indication to exclude structural airway abnormalities and other diagnoses, such as foreign body inhalation. If a clear structural airway abnormality was identified, patients were excluded from the present study. Bronchoscopy was performed under general anesthesia, with a 3.5-mm rigid bronchoscope (Karl Storz GmbH & Co, Tuttingen, Germany), and up to two biopsies were taken from the main carina using biopsy forceps No. 10378L (Karl Storz GmbH&Co).

Carinal biopsies were processed and fixed in formalin before paraffin embedding, microscopic slide preparation and staining. 26 additional biopsy samples were homogenized in Eurozol (EuroClone, Siziano, Italy) or TRIsure (Bioline, London, UK) using the Ultra-Turrax T10 (IKA Labortechnik, Staufen, Germany) to isolate RNA.

### RNA sequencing and Quality Control

RNA samples were processed using the TruSeq RNA sample preparation kit (Illumina, San Diego, CA, USA). The protocol involved quantification of RNA by capillary electrophoresis and polyA amplification to select RNA over DNA. The cDNA fragment libraries were sequenced on an Illumina HiSeq2500 sequencer (paired end 2x150bp). Trimmed fastQ files were aligned to build b37 of the human reference genome using HISAT15 and gene level quantification was performed by HTSeq16 using Ensembl version 75 as gene annotation database. Quality control (QC) metrics were calculated for the raw sequencing data using FastQC and for the aligned reads using Picard-tools. Four RNA samples that did not pass QC were removed and lowly expressed features were filtered out.

### Differential Expression Analysis

Data was normalized by edgeR using TMM, prior to Voom transformation to prepare the data for linear modeling.(*42*) DE analyses were conducted using linear regression separately for three outcomes: asthma, atopy, and AHR. All analyses were adjusted for age (continuous), sex, and smoking exposure (defined as smoking by ≥ 1 parent). lmFit using the eBayes function in the limma R package was used to compute moderated t-statistics for DE analysis. We assessed significance using Benjamini-Hochberg false discovery rate (FDR) <0.05.

### Overrepresentation and gene set enrichment analysis

We selected a list with all nominal significant (p < 0.05) DE genes (n=404, Table S2) and analyzed this list compared to the background list of all genes in our dataset with ConsensusPathDB (*43*). Next, we performed GSEA with the fgsea package in R for published gene sets associated with asthma (*14*), Type2-inflammation (*15*) and activated eosinophils (*16*). We then performed GSEA with gene sets specific for different epithelial cell subsets and cell differentiation trajectories derived from scRNA-seq of tracheal brushes of healthy preschool children.

### Cell type scoring and marker genes

The single cell dataset (*17*), which comprises brushing samples obtained from healthy and COVID-19 positive neonates up to elderly donors at various locations (nose, bronchi, trachea), was analyzed with scanpy (*44*) version 1.8.1. A subset of the dataset was used comprising tracheal samples of Covid-19 negative children of age 4 or less only. The gene list obtained from bulkRNA-seq was filtered for genes with a log fold change (LFC) > 0.8 and a p-value < 0.05 to generate the pre-asthma gene signature (varying the values for log fold change and p-value cutoffs yielded similar results). The expression of this set of genes was then scored in each single cell using “scanpy.tl.score_genes()”. These scores were then plotted on a UMAP embedding and summarized per cell type by their median.

To obtain marker genes for each cell type in the subsetted single cell dataset, “scanpy.tl.rank_genes_groups()” was used, with the “method” parameter set to “logreg”. This employs a logistic regression approach (*45*) to predict cell types by gene expression, followed by ranking genes according to their coefficients in the decision function (also called scores). Marker genes were computed for all epithelial cell types, with all other epithelial cell types as a background. The top 50 lists of marker genes ranked by score per cell type were used for further GSEA in the bulkRNA-seq data.

To obtain marker gene lists for the basal-to-squamous and basal-to-club differentiation trajectories, we combined cell level annotations of the respective cell types into new groups and performed the same procedure as described above on these new groups.

### Cell type enrichment for disease associated loci

Functional GWAS analysis (fGWAS) (*25*) was applied to identify disease relevant cell types as described in detail in (*26*) (https://github.com/natsuhiko/PHM). The model makes use of full summary statistics from GWAS studies, linking SNPs to genes and captures a general trend between gene expression and disease association of linked loci for each cell type. At the same time, it also corrects for linkage disequilibrium (LD) and other relevant factors. We used full summary statistics obtained from two studies (*9, 10*) that both examined adult-onset as well as childhood-onset asthma. Genomic coordinates were mapped to the hg38 genome build using “LiftOver” (*46*).

### Replication in nasal brush samples

Replication analysis was performed in the PIAMA (*23*) birth cohort at age 16 years and in the children from the ARIA study (*24*). Additional information on these cohorts is provided in the online supplement. From the scRNA-seq dataset, we selected a subset of nasal samples of healthy donors below 4 years of age and computed marker gene signatures of the squamous and club cell differentiation trajectories as described above. We performed GSEA with the fgsea package in R with these two gene signatures.

### Stainings

Paraffin embedded biopsies from the original research cohort (n=36) were available for stainings. The biopsies were stained with Cytokeratin 5 and Cytokeratin 14 (details in online supplement), incubated with fluorescent labels, washed and counterstained with DAPI, mounted in aqueous mounting medium and scanned using the Olympus VS200 ASW slide scanner. The images were analyzed with Fuji for Image J. The blinded and randomized images were scored for the presence of co-expression of KRT5/KRT14 by two independent observers. The association between co-expression and asthma outcome at school-age was calculated with R statistical software.

For RNAScope, stored pediatric nasal samples and adult tracheal samples from the Great Ormond Street Hospital were used. The nasal samples were fixed in paraffin blocks and tracheal samples were embedded in OCT at ≅-60°C and 10µm cryosections cut on to superfrost plus slides. Fixed and dried slides (details in online supplement) were loaded on a Leica BOND RX and stained using the RNAScope LS multiplex fluorescent assay (ACD, Bio-Techne) as per the manual. The probes used were human KRT14 and KRT6A. Slides were scanned on a Hamamatsu Nanozoomer S60 microscope at 40X magnification.

## List of Supplementary Materials

Supplementary Materials and Methods: detailed description of methods used in this study.

Table S1. Histological findings according to asthma outcome at school age.

Tables S2 to S12 are submitted in a separate Excel file.

Table S2. All genes DE in pre-asthma vs. transient wheeze with nominal p-value <0.05.

Table S3. Top 100 genes DE in presence vs. absence of atopy at school age.

Table S4. Top 100 genes DE in presence vs. absence of AHR at school age.

Table S5. Pathways and gene ontology categories that are overrepresented (q-value < 0.005) in pre-asthmatic children vs. transient wheeze.

Table S6. Test statistics from DEA of genes in the keratinization GO set.

Table S7. Test statistics from DEA of genes upregulated in the airway epithelium in persistent asthma.

Table S9. Test statistics from DEA of genes upregulated in Type2-inflammation.

Table S9. Test statistics from DEA of genes upregulated in activated airway eosinophils.

Table S10. Top 50 marker genes for the epithelial cell subsets.

Table S11. GSEA results for cell subset specific gene sets.

Table S12. Top 50 marker genes for the basal to squamous cell and basal to club cell differentiation trajectories.

Fig. S1. Upset plot showing overlap between top 50 marker genes for epithelial cell subsets and UMAPs showing expression of selected genes.

Fig. S2. Boxplots showing expression of signature gene sets from epithelial cell subsets.

Fig. S3. Enrichment plots for trajectory gene signatures in nasal brushes.

Fig. S4. Scores for genes most associated with childhood-onset asthma.

Fig. S5. RNAscope images for KRT6a and KRT14 staining.

References (*47-53*)

## Acknowledgments

We kindly acknowledge the Genome Analysis Facility (GAF), Genomics Coordination Centre (GCC) of the University Medical Center Groningen for the RNA Isolation, sample preparation, sequencing and initial quality control of our samples.

## Funding

ZON-MW VICI grant (projectnumber 848101008) (GHK)

Sigrid Jusélius Foundation (KM, MJM)

Wellcome Trust (WT211276/Z/18/Z) (KBM, SAT)

Wellcome Trust Sanger core grant (WT206194) (KBM, SAT)

Chan Zuckerberg Initiative seed network grant (czf2019-002438) (KBM)

Medical Research Council Clinician Scientist Fellowship (MR/W00111X/1) (MZN) Rosetrees Trust (M899) (MZN)

Action Medical Research (GN2911) (MZN)

## Author contributions

Conceptualization: GHK, MB, MCN, MZN, KBM

Data Curation: ETGK, JPP

Formal Analysis: ETGK, JPP

Investigation: KM, MJM, MZN

Methodology: KM, MJM, MNC, GHK, KBM, JPP, ETGK

Resources: JL, MZN, NS, KM, SAT, MCN

Supervision: MJM, GHK, MCN, KBM

Validation: YC, SB, RCHV, JMV, ETGK, GHK

Visualization: JL, WT, MRJ, NS, AWC, KBW

Writing – Original Draft: ETGK, JPP, MZN, MCN, KBM, GHK

Writing – Review & Editing: all co-authors

## Data and materials availability

The participants of the infant bulkRNA-sequencing study did not give written consent for their data to be shared publicly, so after consultation of the legal department of the Helsinki University Central Hospital we are not allowed to publicly share the data under Finnish law.

The single cell dataset is available at the European Genome–phenome Archive under accession number EGAD00001007718. The dataset can be explored interactively through a web portal (https://covid19cellatlas.org).

The data from the nasal bulkRNA-sequencing PIAMA cohort are available in the European Genome-phenome Archive under accession number EGAS00001006240.

The data from the nasal bulkRNA-sequencing ARIA cohort are available at Synapse (www.synapse.org) under project ID syn20687810.

## References and Notes

1. I. V. Yang, C. A. Lozupone, D. A. Schwartz, The environment, epigenome, and asthma. J. Allergy Clin. Immunol. 140, 14–23 (2017).

2. C. Porsbjerg, E. Melén, L. Lehtimäki, D. Shaw, Asthma. Lancet 401, 858–873 (2023).

3. M. Bonato, M. Tiné, E. Bazzan, D. Biondini, M. Saetta, S. Baraldo, Early Airway Pathological Changes in Children: New Insights into the Natural History of Wheezing. J. Clin. Med. Res. 8, 1180 (2019).

4. J. A. Castro-Rodriguez, S. Saglani, C. E. Rodriguez-Martinez, M. A. Oyarzun, L. Fleming, A. Bush, The relationship between inflammation and remodeling in childhood asthma: A systematic review. Pediatr. Pulmonol. 53, 824–835 (2018).

5. K. Malmström, L. P. Malmberg, R. O’Reilly, H. Lindahl, M. Kajosaari, K. M. Saarinen, S. Saglani, F. L. Jahnsen, A. Bush, T. Haahtela, S. Sarna, A. S. Pelkonen, M. J. Mäkelä, Lung function, airway remodeling, and inflammation in infants: outcome at 8 years. Ann. Allergy Asthma Immunol. 114, 90–96 (2015).

6. R. O’Reilly, N. Ullmann, S. Irving, C. J. Bossley, S. Sonnappa, J. Zhu, T. Oates, W. Banya, P. K. Jeffery, A. Bush, S. Saglani, Increased airway smooth muscle in preschool wheezers who have asthma at school age. J. Allergy Clin. Immunol. 131, 1024–1032, 1032.e1–16 (2013).

7. M. Bonato, E. Bazzan, D. Snijders, M. Tinè, D. Biondini, G. Turato, E. Balestro, A. Papi, M. G. Cosio, A. Barbato, S. Baraldo, M. Saetta, Clinical and Pathologic Factors Predicting Future Asthma in Wheezing Children. A Longitudinal Study. Am. J. Respir. Cell Mol. Biol. 59, 458–466 (2018).

8. G. Lezmi, A. Deschildre, R. Abou Taam, M. Fayon, S. Blanchon, F. Troussier, P. Mallinger, B. Mahut, P. Gosset, J. de Blic, Remodelling and inflammation in preschoolers with severe recurrent wheeze and asthma outcome at school age. Clin. Exp. Allergy 48, 806–813 (2018).

9. M. Pividori, N. Schoettler, D. L. Nicolae, C. Ober, H. K. Im, Shared and distinct genetic risk factors for childhood-onset and adult-onset asthma: genome-wide and transcriptome-wide studies. Lancet Respir Med 7, 509–522 (2019).

10. M. A. R. Ferreira, R. Mathur, J. M. Vonk, A. Szwajda, B. Brumpton, R. Granell, B. K. Brew, V. Ullemar, Y. Lu, Y. Jiang, 23andMe Research Team, eQTLGen Consortium, BIOS Consortium, P. K. E. Magnusson, R. Karlsson, D. A. Hinds, L. Paternoster, G. H. Koppelman, C. Almqvist, Genetic Architectures of Childhood- and Adult-Onset Asthma Are Partly Distinct. Am. J. Hum. Genet. 104, 665–684 (2019).

11. M. A. R. Ferreira, R. Mathur, J. M. Vonk, A. Szwajda, B. Brumpton, R. Granell, B. K. Brew, V. Ullemar, Y. Lu, Y. Jiang, 23andMe Research Team, eQTLGen Consortium, BIOS Consortium, P. K. E. Magnusson, R. Karlsson, D. A. Hinds, L. Paternoster, G. H. Koppelman, C. Almqvist, Genetic Architectures of Childhood- and Adult-Onset Asthma Are Partly Distinct. Am. J. Hum. Genet. 104, 665–684 (2019).

12. M. Pividori, N. Schoettler, D. L. Nicolae, C. Ober, H. K. Im, Shared and distinct genetic risk factors for childhood-onset and adult-onset asthma: genome-wide and transcriptome-wide studies. Lancet Respir Med 7, 509–522 (2019).

13. K. Malmström, J. Lohi, L. P. Malmberg, A. Kotaniemi-Syrjänen, H. Lindahl, S. Sarna, A. S. Pelkonen, M. J. Mäkelä, Airway hyperresponsiveness, remodeling and inflammation in infants with wheeze. Clin. Exp. Allergy 50, 558–566 (2020).

14. Y.-H. Tsai, J. S. Parker, I. V. Yang, S. N. P. Kelada, Meta-analysis of airway epithelium gene expression in asthma. Eur. Respir. J. 51, 1701962 (2018).

15. D. F. Choy, B. Modrek, A. R. Abbas, S. Kummerfeld, H. F. Clark, L. C. Wu, G. Fedorowicz, Z. Modrusan, J. V. Fahy, P. G. Woodruff, J. R. Arron, Gene expression patterns of Th2 inflammation and intercellular communication in asthmatic airways. J. Immunol. 186, 1861– 1869 (2011).

16. S. Esnault, E. A. Kelly, E. A. Schwantes, L. Y. Liu, L. P. DeLain, J. A. Hauer, Y. A. Bochkov, L. C. Denlinger, J. S. Malter, S. K. Mathur, N. N. Jarjour, Identification of genes expressed by human airway eosinophils after an in vivo allergen challenge. PLoS One 8, e67560 (2013).

17. M. Yoshida, K. B. Worlock, N. Huang, R. G. H. Lindeboom, C. R. Butler, N. Kumasaka, C. D. Conde, L. Mamanova, L. Bolt, L. Richardson, K. Polanski, E. Madissoon, J. L. Barnes, J. Allen-Hyttinen, E. Kilich, B. C. Jones, A. de Wilton, A. Wilbrey-Clark, W. Sungnak, J. P. Pett, J. Weller, E. Prigmore, H. Yung, P. Mehta, A. Saleh, A. Saigal, V. Chu, J. M. Cohen, C. Cane, A. Iordanidou, S. Shibuya, A.-K. Reuschl, I. T. Herczeg, A. C. Argento, R. G. Wunderink, S. B. Smith, T. A. Poor, C. A. Gao, J. E. Dematte, NU SCRIPT Study Investigators, G. Reynolds, M. Haniffa, G. S. Bowyer, M. Coates, M. R. Clatworthy, F. J. Calero-Nieto, B. Göttgens, C. O’Callaghan, N. J. Sebire, C. Jolly, P. de Coppi, C. M. Smith, A. V. Misharin, S. M. Janes, S. A. Teichmann, M. Z. Nikolić, K. B. Meyer, Local and systemic responses to SARS-CoV-2 infection in children and adults. Nature 602, 321–327 (2022).

18. D. T. Montoro, A. L. Haber, M. Biton, V. Vinarsky, B. Lin, S. E. Birket, F. Yuan, S. Chen, H. M. Leung, J. Villoria, N. Rogel, G. Burgin, A. M. Tsankov, A. Waghray, M. Slyper, J. Waldman, L. Nguyen, D. Dionne, O. Rozenblatt-Rosen, P. R. Tata, H. Mou, M. Shivaraju, H. Bihler, M. Mense, G. J. Tearney, S. M. Rowe, J. F. Engelhardt, A. Regev, J. Rajagopal, A revised airway epithelial hierarchy includes CFTR-expressing ionocytes. Nature 560, 319–324 (2018).

19. H. M. Rigden, A. Alias, T. Havelock, R. O’Donnell, R. Djukanovic, D. E. Davies, S. J. Wilson, Squamous Metaplasia Is Increased in the Bronchial Epithelium of Smokers with Chronic Obstructive Pulmonary Disease. PLoS One 11, e0156009 (2016).

20. F. A. Wolf, F. K. Hamey, M. Plass, J. Solana, J. S. Dahlin, B. Göttgens, N. Rajewsky, L. Simon, F. J. Theis, PAGA: graph abstraction reconciles clustering with trajectory inference through a topology preserving map of single cells. Genome Biol. 20, 59 (2019).

21. J. Cao, M. Spielmann, X. Qiu, X. Huang, D. M. Ibrahim, A. J. Hill, F. Zhang, S. Mundlos, L. Christiansen, F. J. Steemers, C. Trapnell, J. Shendure, The single-cell transcriptional landscape of mammalian organogenesis. Nature 566, 496–502 (2019).

22. C. Trapnell, D. Cacchiarelli, J. Grimsby, P. Pokharel, S. Li, M. Morse, N. J. Lennon, K. J. Livak, T. S. Mikkelsen, J. L. Rinn, The dynamics and regulators of cell fate decisions are revealed by pseudotemporal ordering of single cells. Nat. Biotechnol. 32, 381–386 (2014).

23. A. H. Wijga, M. Kerkhof, U. Gehring, J. C. de Jongste, D. S. Postma, R. C. Aalberse, A. P. H. Wolse, G. H. Koppelman, L. van Rossem, M. Oldenwening, B. Brunekreef, H. A. Smit, Cohort profile: the prevention and incidence of asthma and mite allergy (PIAMA) birth cohort. Int. J. Epidemiol. 43, 527–535 (2014).

24. A. N. Do, Y. Chun, G. Grishina, A. Grishin, A. J. Rogers, B. A. Raby, S. T. Weiss, A. Vicencio, E. E. Schadt, S. Bunyavanich, Network study of nasal transcriptome profiles reveals master regulator genes of asthma. J. Allergy Clin. Immunol. 147, 879–893 (2021).

25. J. K. Pickrell, Joint analysis of functional genomic data and genome-wide association studies of 18 human traits. Am. J. Hum. Genet. 94, 559–573 (2014).

26. R. Elmentaite, N. Kumasaka, K. Roberts, A. Fleming, E. Dann, H. W. King, V. Kleshchevnikov, M. Dabrowska, S. Pritchard, L. Bolt, S. F. Vieira, L. Mamanova, N. Huang, F. Perrone, I. Goh Kai’En, S. N. Lisgo, M. Katan, S. Leonard, T. R. W. Oliver, C. E. Hook, K. Nayak, L. S. Campos, C. Domínguez Conde, E. Stephenson, J. Engelbert, R. A. Botting, K. Polanski, S. van Dongen, M. Patel, M. D. Morgan, J. C. Marioni, O. A. Bayraktar, K. B. Meyer, X. He, R. A. Barker, H. H. Uhlig, K. T. Mahbubani, K. Saeb-Parsy, M. Zilbauer, M. R. Clatworthy, M. Haniffa, K. R. James, S. A. Teichmann, Cells of the human intestinal tract mapped across space and time. Nature 597, 250–255 (2021).

27. A. Bush, A. Menzies-Gow, Phenotypic differences between pediatric and adult asthma. Proc. Am. Thorac. Soc. 6, 712–719 (2009).

28. Y. Zhang, K. E. Black, T.-K. N. Phung, S. R. Thundivalappil, T. Lin, W. Wang, J. Xu, C. Zhang, L. P. Hariri, A. Lapey, H. Li, P. H. Lerou, X. Ai, J. Que, J.-A. Park, B. P. Hurley, H. Mou, Human Airway Basal Cells Undergo Reversible Squamous Differentiation and Reshape Innate Immunity. Am. J. Respir. Cell Mol. Biol. 68, 664–678 (2023).

29. M. A. Ferreira, J. M. Vonk, H. Baurecht, I. Marenholz, C. Tian, J. D. Hoffman, Q. Helmer, A. Tillander, V. Ullemar, J. van Dongen, Y. Lu, F. Rüschendorf, J. Esparza-Gordillo, C. W. Medway, E. Mountjoy, K. Burrows, O. Hummel, S. Grosche, B. M. Brumpton, J. S. Witte, J.-J. Hottenga, G. Willemsen, J. Zheng, E. Rodríguez, M. Hotze, A. Franke, J. A. Revez, J. Beesley, M. C. Matheson, S. C. Dharmage, L. M. Bain, L. G. Fritsche, M. E. Gabrielsen, B. Balliu, 23andMe Research Team, AAGC collaborators, BIOS consortium, LifeLines Cohort Study, J. B. Nielsen, W. Zhou, K. Hveem, A. Langhammer, O. L. Holmen, M. Løset, G. R. Abecasis, C. J. Willer, A. Arnold, G. Homuth, C. O. Schmidt, P. J. Thompson, N. G. Martin, D. L. Duffy, N. Novak, H. Schulz, S. Karrasch, C. Gieger, K. Strauch, R. B. Melles, D. A. Hinds, N. Hübner, S. Weidinger, P. K. E. Magnusson, R. Jansen, E. Jorgenson, Y.-A. Lee, D. I. Boomsma, C. Almqvist, R. Karlsson, G. H. Koppelman, L. Paternoster, Shared genetic origin of asthma, hay fever and eczema elucidates allergic disease biology. Nat. Genet. 49, 1752–1757 (2017).

30. M. A. R. Ferreira, J. M. Vonk, H. Baurecht, I. Marenholz, C. Tian, J. D. Hoffman, Q. Helmer, A. Tillander, V. Ullemar, Y. Lu, S. Grosche, F. Rüschendorf, R. Granell, B. M. Brumpton, L. G. Fritsche, L. Bhatta, M. E. Gabrielsen, J. B. Nielsen, W. Zhou, K. Hveem, A. Langhammer, O. L. Holmen, M. Løset, G. R. Abecasis, C. J. Willer, N. C. Emami, T. B. Cavazos, J. S. Witte, A. Szwajda, 23andMe Research Team, collaborators of the SHARE study, D. A. Hinds, N. Hübner, S. Weidinger, P. K. Magnusson, E. Jorgenson, R. Karlsson, L. Paternoster, D. I. Boomsma, C. Almqvist, Y.-A. Lee, G. H. Koppelman, Age-of-onset information helps identify 76 genetic variants associated with allergic disease. PLoS Genet. 16, e1008725 (2020).

31. L. Möbus, E. Rodriguez, I. Harder, D. Stölzl, N. Boraczynski, S. Gerdes, A. Kleinheinz, S. Abraham, A. Heratizadeh, C. Handrick, E. Haufe, T. Werfel, J. Schmitt, S. Weidinger, TREATgermany study group, Atopic dermatitis displays stable and dynamic skin transcriptome signatures. J. Allergy Clin. Immunol. 147, 213–223 (2021).

32. D. H. Kim, A. Gu, J.-S. Lee, E. J. Yang, A. Kashif, M. H. Hong, G. Kim, B. S. Park, S. J. Lee, I. S. Kim, Suppressive effects of S100A8 and S100A9 on neutrophil apoptosis by cytokine release of human bronchial epithelial cells in asthma. Int. J. Med. Sci. 17, 498–509 (2020).

33. N. Lopez-Souza, G. Dolganov, R. Dubin, L. A. Sachs, L. Sassina, H. Sporer, S. Yagi, D. Schnurr, H. A. Boushey, J. H. Widdicombe, Resistance of differentiated human airway epithelium to infection by rhinovirus. Am. J. Physiol. Lung Cell. Mol. Physiol. 286, L373–81 (2004).

34. S. Schloer, D. Pajonczyk, U. Rescher, Annexins in Translational Research: Hidden Treasures to Be Found. Int. J. Mol. Sci. 19, 1781 (2018).

35. R. Granell, J. A. Curtin, S. Haider, N. T. Kitaba, S. A. Mathie, L. G. Gregory, L. L. Yates, M. Tutino, J. Hankinson, M. Perretti, J. M. Vonk, H. S. Arshad, P. Cullinan, S. Fontanella, G. C. Roberts, G. H. Koppelman, A. Simpson, S. W. Turner, C. S. Murray, C. M. Lloyd, J. W. Holloway, A. Custovic, UNICORN and Breathing Together Investigators, A meta-analysis of genome-wide association studies of childhood wheezing phenotypes identifies ANXA1 as a susceptibility locus for persistent wheezing. Elife 12, e84315 (2023).

36. S. T. Holgate, D. E. Davies, P. M. Lackie, S. J. Wilson, S. M. Puddicombe, J. L. Lordan, Epithelial-mesenchymal interactions in the pathogenesis of asthma. J. Allergy Clin. Immunol. 105, 193–204 (2000).

37. A. Kaplan, M. Bueno, A. E. Fournier, Extracellular functions of 14-3-3 adaptor proteins. Cell. Signal. 31, 26–30 (2017).

38. K. Rietscher, R. Keil, A. Jordan, M. Hatzfeld, 14-3-3 proteins regulate desmosomal adhesion via plakophilins. J. Cell Sci. 131, jcs212191 (2018).

39. N. Asdaghi, R. T. Kilani, A. Hosseini-Tabatabaei, S. O. Odemuyiwa, T.-L. Hackett, D. A. Knight, A. Ghahary, R. Moqbel, Extracellular 14-3-3 from human lung epithelial cells enhances MMP-1 expression. Mol. Cell. Biochem. 360, 261–270 (2012).

40. R. Rajah, R. V. Nachajon, M. H. Collins, H. Hakonarson, M. M. Grunstein, P. Cohen, Elevated levels of the IGF-binding protein protease MMP-1 in asthmatic airway smooth muscle. Am. J. Respir. Cell Mol. Biol. 20, 199–208 (1999).

41. M. Deprez, L.-E. Zaragosi, M. Truchi, C. Becavin, S. Ruiz García, M.-J. Arguel, M. Plaisant, V. Magnone, K. Lebrigand, S. Abelanet, F. Brau, A. Paquet, D. Pe’er, C.-H. Marquette, S. Leroy, P. Barbry, A Single-Cell Atlas of the Human Healthy Airways. Am. J. Respir. Crit. Care Med. 202, 1636–1645 (2020).

42. C. W. Law, Y. Chen, W. Shi, G. K. Smyth, voom: Precision weights unlock linear model analysis tools for RNA-seq read counts. Genome Biol. 15, R29 (2014).

43. R. Herwig, C. Hardt, M. Lienhard, A. Kamburov, Analyzing and interpreting genome data at the network level with ConsensusPathDB. Nat. Protoc. 11, 1889–1907 (2016).

44. F. A. Wolf, P. Angerer, F. J. Theis, SCANPY: large-scale single-cell gene expression data analysis. Genome Biol. 19, 15 (2018).

45. V. Ntranos, L. Yi, P. Melsted, L. Pachter, A discriminative learning approach to differential expression analysis for single-cell RNA-seq, Nat. Methods. 16, 163–166 (2019).

46. A. S. Hinrichs, D. Karolchik, R. Baertsch, G. P. Barber, G. Bejerano, H. Clawson, M. Diekhans, T. S. Furey, R. A. Harte, F. Hsu, J. Hillman-Jackson, R. M. Kuhn, J. S. Pedersen, A. Pohl, B. J. Raney, K. R. Rosenbloom, A. Siepel, K. E. Smith, C. W. Sugnet, A. Sultan-Qurraie, D. J. Thomas, H. Trumbower, R. J. Weber, M. Weirauch, A. S. Zweig, D. Haussler, W. J. Kent, The UCSC Genome Browser Database: update 2006. Nucleic Acids Res. 34, D590–8 (2006).

47. L. P. Malmberg, K. Malmström, A. Kotaniemi-Syrjänen, J. Lohi, A. S. Pelkonen, S. Sarna, M. J. Mäkelä, Early bronchial inflammation and remodeling and airway hyperresponsiveness at school age. Allergy 75, 1765–1768 (2020).

48. D. Kim, B. Langmead, S. L. Salzberg, HISAT: a fast spliced aligner with low memory requirements. Nat. Methods 12, 357–360 (2015).

49. H. Li, B. Handsaker, A. Wysoker, T. Fennell, J. Ruan, N. Homer, G. Marth, G. Abecasis, R. Durbin, 1000 Genome Project Data Processing Subgroup, The Sequence Alignment/Map format and SAMtools. Bioinformatics 25, 2078–2079 (2009).

50. S. Anders, P. T. Pyl, W. Huber, HTSeq--a Python framework to work with high-throughput sequencing data. Bioinformatics 31, 166–169 (2015).

51. C. W. Law, M. Alhamdoosh, S. Su, X. Dong, L. Tian, G. K. Smyth, M. E. Ritchie, RNA-seq analysis is easy as 1-2-3 with limma, Glimma and edgeR. F1000Res. **5** (2016).

52. C. Qi, Y. Jiang, I. V. Yang, E. Forno, T. Wang, J. M. Vonk, U. Gehring, H. A. Smit, E. B. Milanzi, O. A. Carpaij, M. Berg, L. Hesse, S. Brouwer, J. Cardwell, C. J. Vermeulen, E. Acosta-Pérez, G. Canino, N. Boutaoui, M. van den Berge, S. A. Teichmann, M. C. Nawijn, W. Chen, J. C. Celedón, C.-J. Xu, G. H. Koppelman, Nasal DNA methylation profiling of asthma and rhinitis. J. Allergy Clin. Immunol. 145, 1655–1663 (2020).

53. W. C. Moore, E. R. Bleecker, D. Curran-Everett, S. C. Erzurum, B. T. Ameredes, L. Bacharier, W. J. Calhoun, M. Castro, K. F. Chung, M. P. Clark, R. A. Dweik, A. M. Fitzpatrick, B. Gaston, M. Hew, I. Hussain, N. N. Jarjour, E. Israel, B. D. Levy, J. R. Murphy, S. P. Peters, W. G. Teague, D. A. Meyers, W. W. Busse, S. E. Wenzel, National Heart, Lung, Blood Institute’s Severe Asthma Research Program, Characterization of the severe asthma phenotype by the National Heart, Lung, and Blood Institute’s Severe Asthma Research Program. J. Allergy Clin. Immunol. 119, 405–413 (2007).

