## Supplementary materials and methods for "Childhood-onset asthma is characterized by airway epithelial hillock-to-squamous differentiation in early life"

**Study Population**

Full-term infants who had been referred to Helsinki University Central Hospital, a tertiary center, for investigation of recurrent respiratory symptoms, including dyspnea, cough, and wheeze, and were symptomatic for at least 4 weeks, were included. As part of their clinical work-up, all infants with a history of physician-diagnosed wheeze and either persistent (daily to weekly) or recurrent troublesome breathing were studied using rigid bronchoscopy. Patients were excluded if they had a need for inhaled corticosteroids (ICS) within 8 weeks prior to the first visit; a cumulative life-time systemic prednisolone use for more than 3 days at a dose of 2 mg/kg, or an equipotent dose of another systemic corticosteroid or life-time ICS use more than 4 weeks; respiratory infection in the 14 days preceding the lung function measurement; any obvious structural defect; any suspicion of ciliary dyskinesia, cystic fibrosis or immune deficiencies. Other exclusion criteria were: prematurity (<36/40 weeks gestation) or small for gestational age (birth weight < -2 SD); height at the time of investigation < -2.5 SD; bronchopulmonary dysplasia; and specific airways conductance (sGaw) <35% of predicted combined with dyspnea at rest.

Chest X-ray and skin-prick tests were performed. Blood specimens were obtained for analysis of blood eosinophil count and serum total immunoglobulin E (IgE) levels a.o.

**Follow up at school age**

Lung function and bronchodilator responsiveness were measured using a pneumotachograph-based flow-volume spirometer (Masterscreen Pneumo, Jaeger GmbH, Wurzburg, Germany). Airway hyperresponsiveness (AHR) was evaluated with a dosimetric  methacholine challenge, as previously described *(47)**.* A calibrated  nebulizer (Salter Labs 8900, Arvin, CA) was connected to an automatic, inhalation-synchronized dosimeter (Spira Electro II, Spira Respiratory Care Center Ltd, Finland). By calculating the number of breaths with nebulized methacholine, a dosage scheme with four non-cumulative dose steps was delivered (0.1, 0.3, 0.9 and 1.8  mg), with V’maxFRC being recorded after each dose. There were two end-points in the challenge test; a fall of 40% or more in V’maxFRC or reaching the maximal dose of methacholine. The provocative dose of methacholine causing a 40% fall in V’maxFRC (PD40 V’maxFRC) was determined from the dose-response curves. Oxygen saturation and heart rate were continuously monitored with a pulse oximeter (Biox 3700e, Ohmeda, Louisville, KY).

**Bronchoscopy and biopsy processing**

Bronchoscopy was performed to exclude structural airway abnormalities, such as subglottic stenosis, laryngo-, tracheo-, or bronchomalacia, and other diagnoses, such as foreign body inhalation or mucus plugging.

Carinal biopsies were processed and fixed in formalin before paraffin embedding,

microscopic slide preparation and staining *(13)**.* Thickness of the reticular basement membrane (RBM) was estimated by grid-overlay method, as previously described *(13)**.* The amount of smooth muscle was expressed as a percentage of subepithelial area of the biopsy (ASM%) *(13)*.

Inflammatory cells were identified in mucosa and submucosa by immunostaining using antibodies: T-lymphocytes (CD3, 2GVG Ventana, Roche), B-lymphocytes (CD20, L26 Ventana, Roche), plasma cells (CD138, B-A38, Ventana, Roche), mast cells (CD117, polyclonal, Dako), and macrophages (CD163, 10D6, Novocastra).  Results were expressed as number of cells/subepithelial area (1/mm^2^).

Eosinophils were counted from hematoxylin-eosin slides. Neutrophilic leukocytes and eosinophils were stained with CD15 (MMA, Roche) and identified based on morphology. Results were expressed as number of cells/subepithelial area (1/mm^2^).

**RNA isolation, sample preparation and sequencing**

For RNA isolation, samples were homogenized in Eurozol (EuroClone, Siziano, Italy) or TRIsure (Bioline, London, UK) using the Ultra-Turrax T10 (IKA Labortechnik, Staufen, Germany). Glycogen (Roche Diagnostics GmbH, Mannheim, Germany) was used as a carrier molecule during RNA isolation (20 µg / sample). Total RNA extraction was performed according to the manufacturer’s instructions (Eurozol / TRIsure), and RNA was dissolved in 25 µl of diethylpyrocarbonate (DEPC) treated water and stored at -70°C.

Initial quality control (QC) and RNA quantification of the samples was performed by capillary electrophoresis using the LabChip GX (Perkin Elmer, Waltham, MA, USA). PolyA amplification was used to select RNA over DNA and account for DNA contamination that was found during initial QC.

Sequence libraries were generated using the TruSeq RNA sample preparation kits (Illumina, San Diego, CA, USA) using the Sciclone NGS Liquid Handler (Perkin Elmer). In case of contamination of adapter duplexes an extra purification of the libraries was performed with the automated agarose gel separation system Labchip XT (Perkin Elmer). The obtained cDNA fragment libraries were sequenced on an Illumina HiSeq2500 using default parameters (paired end 2x150bp) in pools of multiple samples.

The RNA Integrity (RIN) Scores and RNA-concentrations (in ng / μL) were analyzed. When RNA concentrations are very low, RIN-scores as called by the bioanalyzer are unreliable or absent. In this case, the presence of 18S/28S ribosomal fragment peaks on visual inspection was used as a substitute. Two samples were excluded based on a low RNA-concentration (in ng / μL) in combination with an absent 18S/28S ribosomal fragment peak on visual inspection of the gel electropherogram.

**Gene expression quantification**

The trimmed fastQ files were aligned to build b37 human reference genome using HISAT (version 0.1.5) *(48)* allowing for 2 mismatches. Of the 1.69E^9^ raw reads, 1.21E^9^ could be aligned. Before gene quantification SAMtools (version 1.2) *(49)* was used to sort the aligned reads.

The gene level quantification was performed by HTSeq (version 0.6.1p1) *(50)* using --mode=union--stranded=no and, Ensembl version 75 was used as gene annotation database.

**Quality Control**

QC metrics were calculated for the raw sequencing data. This was done using the tool FastQC (FastQC/0.11.3-Java-1.7.0_80). With FastQC, several aspects of read quality are evaluated in different analysis modules and samples get a PASS, WARN or FAIL flag for each of those modules. This gives a rough indication of sequencing quality, but very poor samples tend to stand out by failing in multiple modules.

The quality of the alignment was assessed using five metrics of Picard tools (Picard-tools picard/1.130-Java-1.7.0_80): proportion of high quality of aligned reads, proportion of paired reads, 3’ to 5’ bias, proportion of ribosomal bases and insert size. We enforced a threshold of >0.60 proportion high quality mapping reads and >0.85 proportion of aligned reads. For the 3’ to 5’ bias, we did not enforce a threshold, as the poly-A protocol is sensitive to provoke 3’ bias in this metric. For the proportion of ribosomal bases we enforced a threshold of <5%. The insert size corresponded to the selected fragment size during sample preparation, and no samples were extremely deviant from the mean.

**Principal Component Analysis**

The first three principal components were analyzed based on the normalized counts in two dimensional graphs. Two samples that failed previous QC steps also appeared to be outliers on the PCA. These samples were excluded from further analysis. Further PCA analysis showed no clustering based on well or year of biopsy. We performed a gender check in which a PCA and clustering were done using only X and Y genes to check if samples had been connected to the right phenotype data. All samples were marked with the correct gender.

**Differential Expression Analysis**

Samples that did not pass QC were removed. Lowly expressed features were filtered out, by removing all features with <5/M (M = median library size in millions) counts per million in at least half the samples. Out of 38,608 genes, 20,294 genes are kept after this filtering method.

Differential expression analysis (DEA) was performed in R. Data was normalized by edgeR using TMM. The R package Limma using the function Voom *(42)* was used to transform count data to logCPM (counts per million) to prepare the normalized data for linear modeling *(51)**.* A design model was built using gender, age, smoke exposure and current asthma as coefficients.

A linear model fit was made using the voom matrix and the design matrix containing the coefficient data. The function lmFit in R using the eBayes function was used to compute moderated t-statistics for DEA. A top list of genes which were differentially expressed with the lowest p-value was generated with the R command topTable. DEA was repeated for presence or absence of atopy and AHR at school age.

**Additional analysis with GWAS SNPs**

For the fGWAS analysis summary statistics were obtained from two studies *(9, 10)* that both examined adult-onset as well as childhood-onset asthma. For the study by Ferreira et al. *(10)* summary statistics were obtained from the EBI GWAS catalog (<https://www.ebi.ac.uk/gwas/studies/GCST007800>) and for the study by Pividori et al. *(9)* from zenodo (<https://zenodo.org/record/3248979#.YWAvub8o9hE>).

We analyzed the contribution of genes with the largest disease association. To this end we selected the top 500 genes associated with childhood-onset asthma from each GWAS study and used the intersection (415 genes) for analysis of their expression across cells in the single cell reference (Fig. S4 A+B) as done previously for the pre-asthma gene signature.

**Nasal brush replication**

*PIAMA birth cohort*

The first replication analysis was performed in the PIAMA (Prevention and Incidence of Asthma and Mite Allergy) birth cohort at age 16 years. Details of the cohort have been published previously *(23)**.* The study started with 3,963 newborns. Questionnaire based follow-up of the children took place at 3 months of age, annually from 1 to 8 years of age, and at 11, 14, and 16 years of age, with clinical investigations at ages 4, 8, 12 and 16 years. The Medical Ethical Committees of the participating institutes approved the study and the parents and legal guardians of all participants as well as the participants themselves gave written informed consent.

The current study used the nasal brush samples from the 16 year visit (mean age 16.3 ± 0.2 years). The cohort consisted of 56.7% females and ~97% of children had European white ancestry. Brushing was performed with Cytosoft brush CP-5B (Cyto-Pak) after local anesthesia with 1% lidocaine spray, from the lateral area underneath the inferior turbinate. RNA was extracted from the nasal brushes and sequencing was performed on Illumina HiSeq2500 platform as previously described *(52)**.*

QC metrics were calculated for the raw sequencing data, using the FastQC tool (version 0.11.3). Alignments of RNA of 333 subjects were obtained. QC metrics were calculated for the aligned reads using Picard-tools (version 1.130) CollectRnaSeqMetrics, MarkDuplicates, CollectInsertSize-Metrics and SAMtools flagstat. After QC, 326 subjects and 17,156 genes were retained. Raw count data were transformed to log2CPM using voom and analyzed in the limma package.

Asthma was defined as the presence of at least 2 out of the following 3 criteria: 1) Doctor diagnosed asthma ever; 2) Wheeze in the last 12 months; and 3) Prescription of asthma medication in the last 12 months.

**ARIA cohort**

Children from the AiRway In Asthma study (ARIA) *(24)* served as a second validation cohort. These participants included 156 children with mean age 13.1 years (SD=3.7), of which 34 children had severe persistent asthma and 122 were healthy controls. ARIA participants were recruited from the Mount Sinai Health System in the New York metropolitan area, USA, with written informed consent and Mount Sinai IRB approval. The cohort was balanced with regard to sex (51.3% female), and race/ethnicity (12.8% Black, 23.7% Latino, 46.2% White, 17.3% Asian and other). Children with asthma had their severe persistent asthma diagnosed by a pediatric pulmonologist based on the Severe Asthma Research Program criteria *(53)*; 26.5% had been hospitalized and 61.8% had had an emergency department visit for asthma in the past year *(24)**.* Healthy controls had no personal or family history of asthma and demonstrated normal spirometry results with no bronchodilator response. Nasal brushing, RNA isolations, RNA sequencing, mapping, and quality control were performed as previously described *(24)**.*

**Stainings**

Paraffin embedded biopsies from 36 participants of the original research cohort were available for stainings. These were cut to 4 µm slides, deparaffinized and boiled in Tris-EDTA buffer in a microwave as antigen-retrieval method. The biopsies were stained with rabbit-anti-human Cytokeratin 5 (Abcam ab52635), 1/800 and mouse-anti-human Cytokeratin 14 (Sino Biological 100129-MM01T), 1/10,000, for 1 hour (RT). After washing, the slides were incubated with donkey-anti-mouse Alexa Fluor 647 (Invitrogen A31517), 1/200, and donkey-anti-rabbit Alexa Fluor 555 (Invitrogen A31572), 1/200, for 30 minutes (RT). After washing, the slides were counterstained with 1/500 diluted DAPI, mounted in aqueous mounting medium and scanned using the Olympus VS200 ASW slide scanner. The images were analyzed with Fuji for Image J. With a homemade macro the captured gray-scale images were converted into colored images, stacked and saved as a tiff file. The blinded and randomized images were scored for the total abundance of co-expression of Cytokeratin 5 and 14 by two independent observers. Per patient the antibody stained biopsy was compared with the image of a biopsy that was only incubated with both secondary antibodies and stained with DAPI.

For RNAScope, stored pediatric nasal samples and adult tracheal samples from the Great Ormond Street Hospital were used. Ethical approval was given through REC reference 20/PR/0542, IRAS project ID 253088, London - Bromley Research Ethics Committee, administered through the Great Ormond Street Hospital NHS Foundation Trust.

The nasal samples were fixed in paraffin blocks and tracheal samples were embedded in OCT at ≅-60°C and 10µm cryosections cut on to superfrost plus slides. Immediately prior to RNAScope, slides were fixed for 15 minutes with 4°C PFA, followed by 90 minutes in room temperature PFA and dehydrated through a standard ethanol series (50%, 70%, 100%, 100%). Dried slides were loaded on a Leica BOND RX and stained using the RNAScope LS multiplex fluorescent assay (ACD, Bio-Techne) as per the manual, with no epitope retrieval, protease IV pre-treatment for 30min at room temperature, opal dyes 520 /570 / 650 at 1:1000 dilution and DAPI at 1:50,000 (Invitrogen D1306, 5mg/ml stock).  The probes used were ACD (Bio-Techne) human KRT14 (310198), KRT6A (520728-C2) and LY6D (484688-C3).  Slides were scanned on a Hamamatsu Nanozoomer S60 microscope at 40X magnification.

**Table S1. Histological findings according to asthma outcome at school age**

|  | **Asthma (n = 10)** | **Transient wheeze (n = 12)** |
| --- | --- | --- |
| RBM thickness (µm) | 3.8 (3.1 – 4.9) | 4.1 (3.1 – 5.2) |
| ASM / subepithelial area (%) | 0.10 (0 – 0.22) | 0.11 (0.01 – 0.23) |
| Mast cells / subepithelial area  (no. per 0.1mm^2^) | 153.5 (0 – 269.2) | 112.0 (71.3 – 225.7) |
| Mast cells / ASM  (no. per 0.1mm^2^) | 28.9 (0 – 104.1) | 39.9 (0 – 87.7) |
| Eosinophils / subepithelial area  (no. per 0.1mm^2^) | 0.82 (0 – 188.2) | 0 (0 – 3.8) |
| Neutrophils / subepithelial area  (no. per 0.1mm^2^) | 118.1 (35.6 – 466.3) | 72.0 (28.4 – 332.0) |

Legend: Median (range) of reticular basement membrane (RBM) thickness, proportion of airway smooth muscle (ASM) in the subepithelial area and counts (per 0.1mm^2^) of mast cells, eosinophils and neutrophils in the subepithelial area and counts of mast cells in the ASM layer. None of these outcomes was significantly different.

**Tables S2 to S12 are submitted in a separate Excel file.**

**Figure legends**

**Fig. S1. (A)** Upset plot showing overlap between top 50 marker genes selected for the 14 epithelial cell clusters.

**(B)** UMAPs showing expression of selected genes.

**(C)** Descending pointplots showing gene expression of ANXA1 and ANXA2 per cell type.

**Fig. S2. Boxplots showing expression of signature gene sets from epithelial cell subsets.** Median and 25-75 IQR of expression (average log2 counts per million) of signature gene sets from all 14 epithelial cell subsets in children <2 years in relation to asthma at school age.

**Fig. S3. Enrichment plots for trajectory gene signatures in nasal brushes.**

Normalized enrichment scores with adjusted p-value of squamous **(A, C)** and club **(B, D)** cell differentiation trajectory gene signatures in nasal brushes from adolescents in relation to asthma status in the PIAMA cohort **(A, B)** and ARIA cohort **(C, D)**.

**Fig. S4. Scores for genes most associated with childhood-onset asthma.**

**(A)** Scores for a set of 415 common genes most associated with childhood-onset asthma across both GWAS studies were computed based on their expression per cell. Cell subsets are ranked by the median score (points). 95% confidence intervals (lines) are shown per cell subset.

**(B)** Score on UMAP.

**(C)** Heatmap of gene expression for 30 genes out of 415 childhood-onset asthma associated genes that are most highly expressed across cell subsets.

**Fig. S5. RNAscope images for KRT6a and KRT14 staining.**

**(A**) PFA fixed and paraffin embedded pharyngeal sections from a 1.5 year old healthy child stained with KRT6a and KRT14 RNAscope probes. Scale bar 50µm

**(B/C)** Fresh frozen tracheal sections from adult airway from a deceased organ donor stained with KRT6a and KRT14 RNAscope probes. Scale bar 100µm (with H&E).
