## Supplementary figures and images for "Childhood-onset asthma is characterized by airway epithelial hillock-to-squamous differentiation in early life"

### Fig. S1

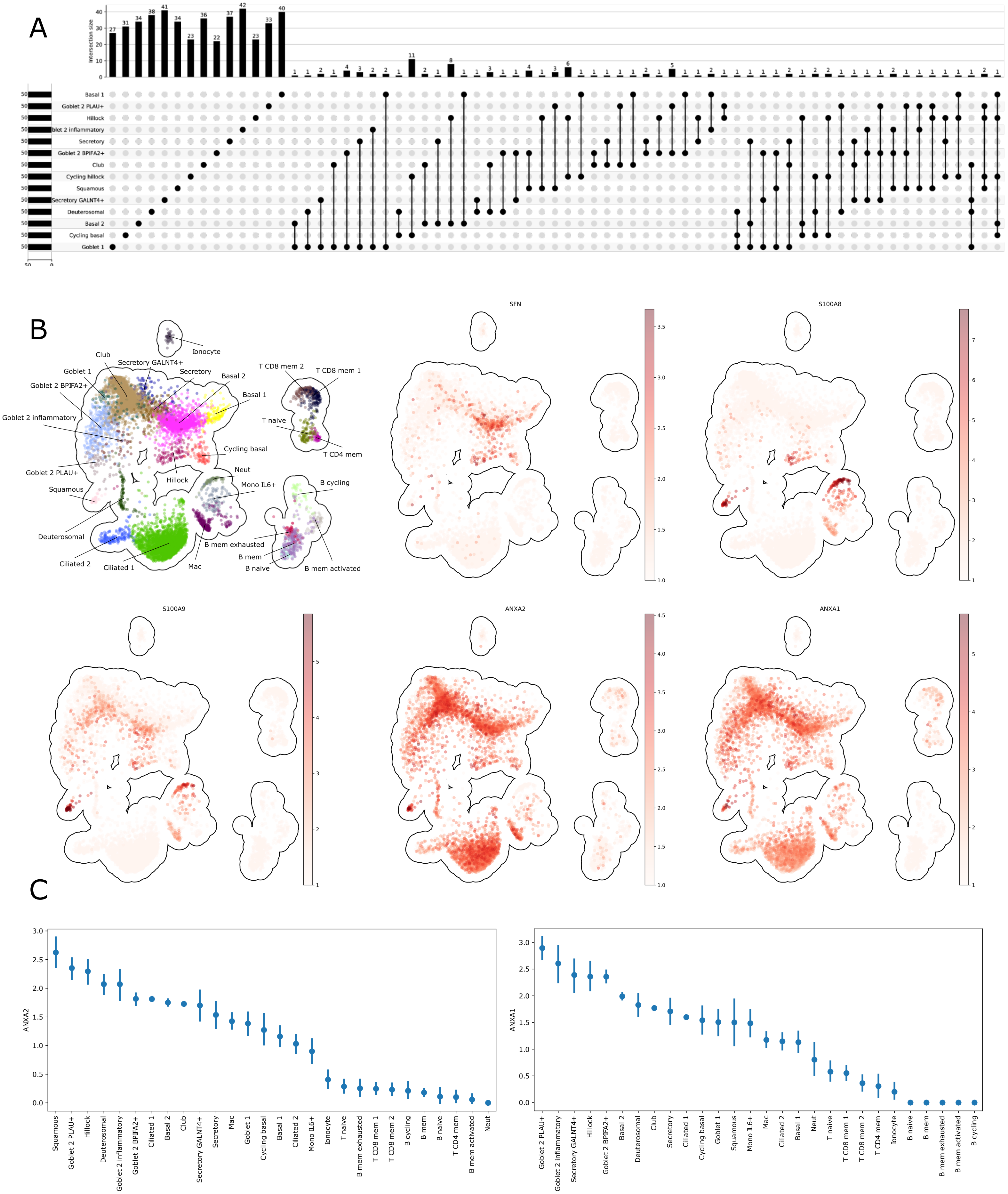

### Fig. S2

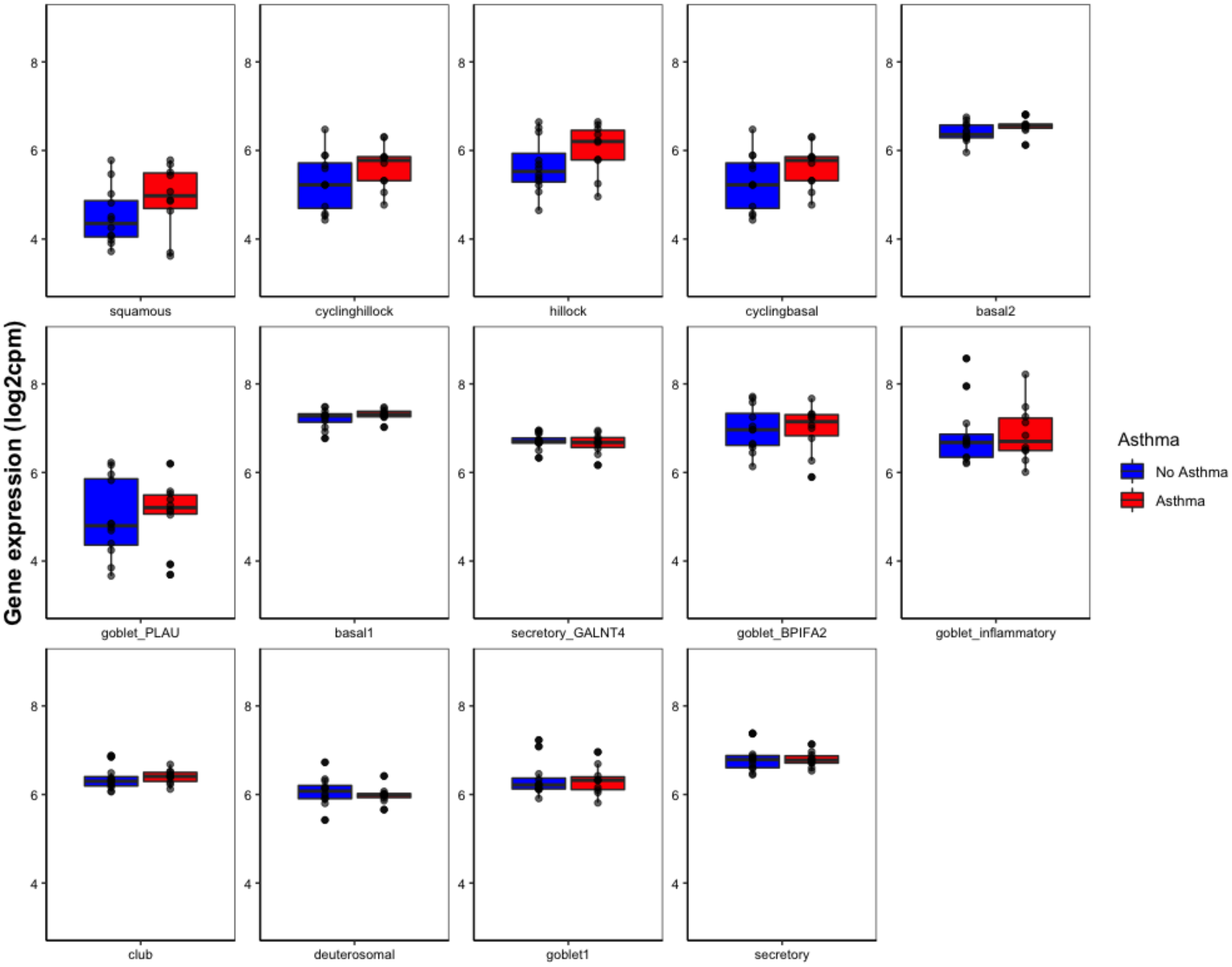

### Fig. S3

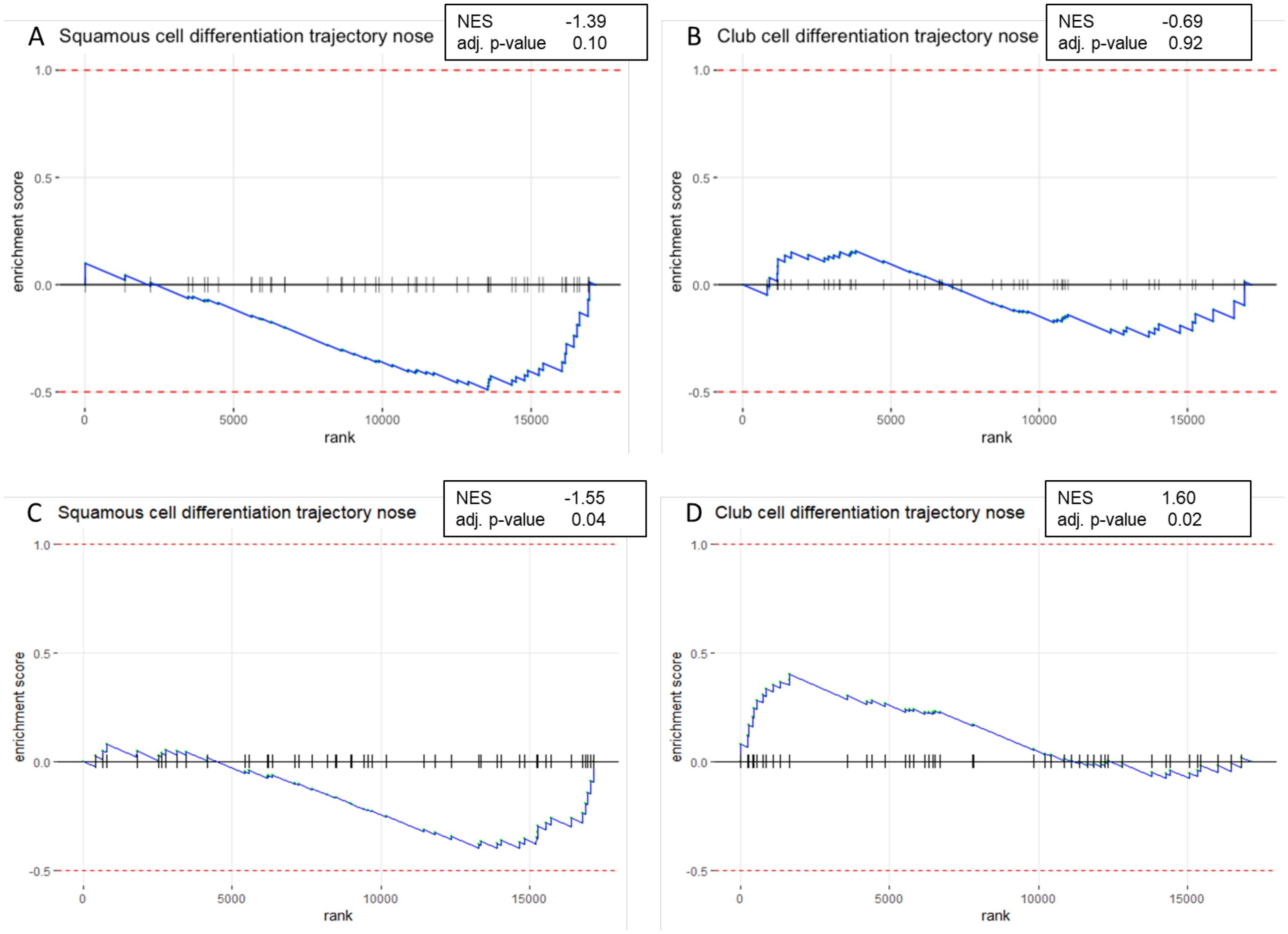

### Fig. S4

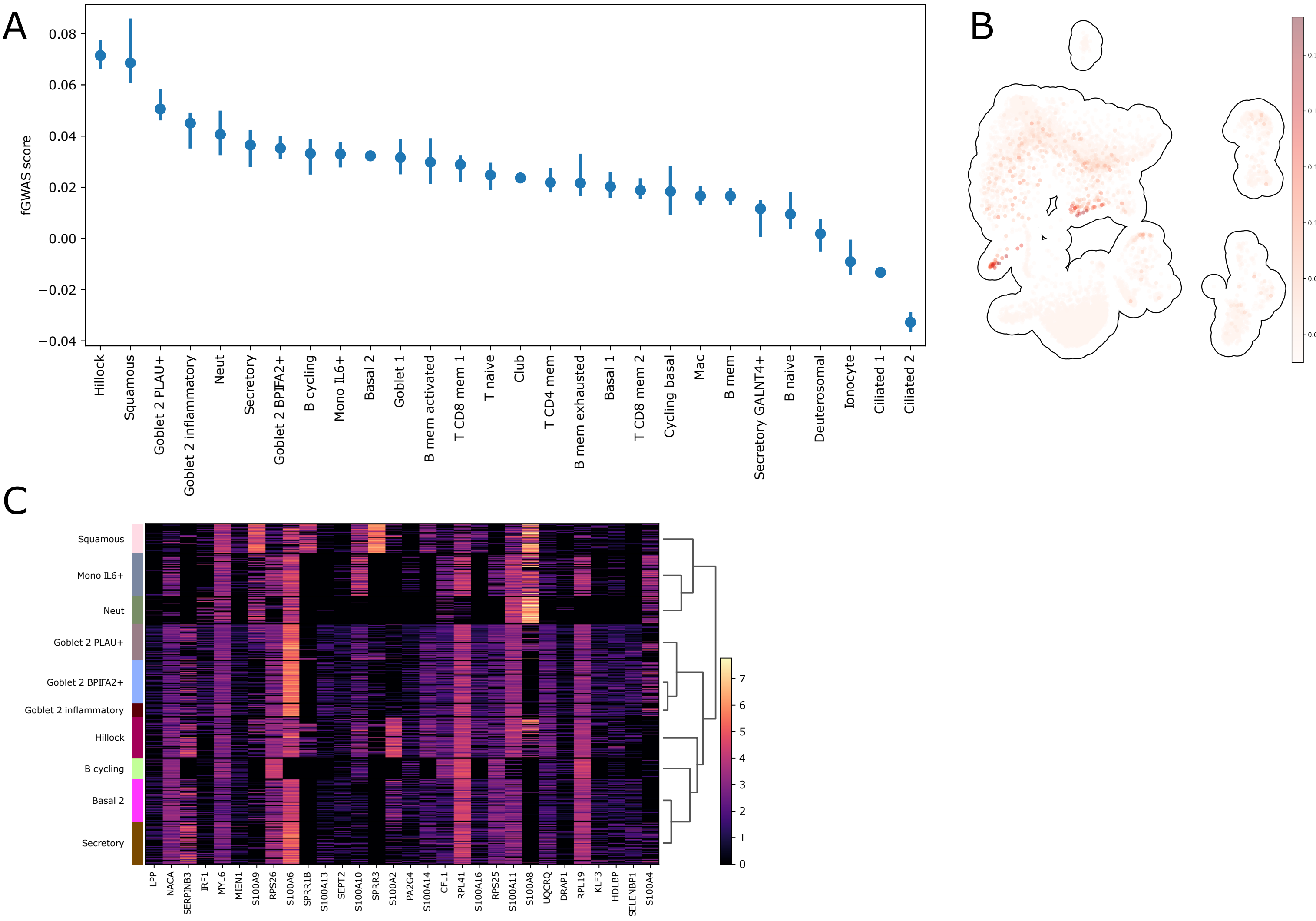

### Fig. S5

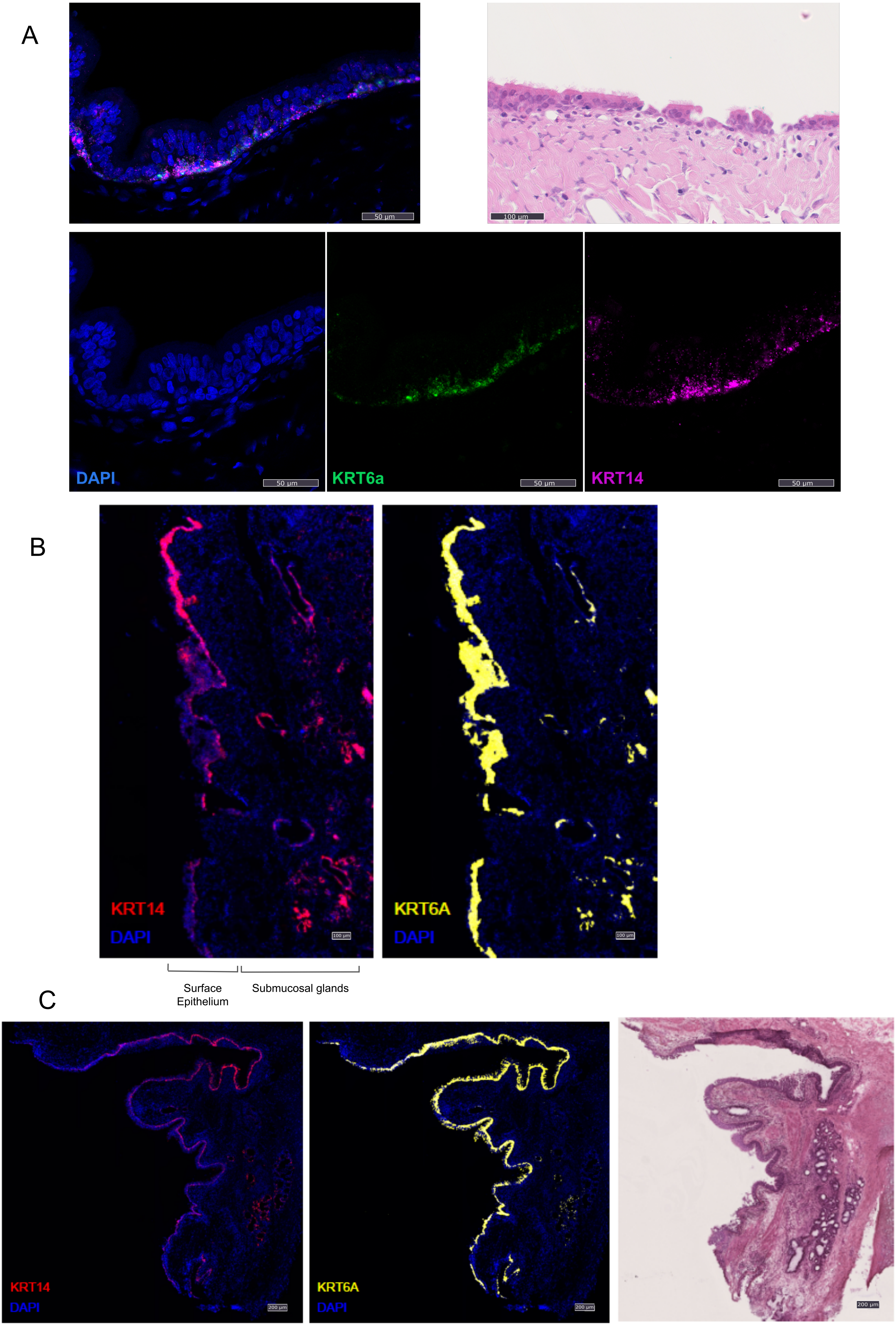
